# The Molecular Logic Organizing the Functional Compartmentalization of Reciprocal Synapses

**DOI:** 10.1101/2021.05.24.445461

**Authors:** Cosmos Yuqi Wang, Justin H. Trotter, Kif Liakath-Ali, Sung-Jin Lee, Xinran Liu, Thomas C. Südhof

**Affiliations:** Dept. of Molecular and Cellular Physiology, School of Medicine, Stanford University, Stanford, CA 94305, USA; Dept. of Cell Biology, School of Medicine, Yale University, New Haven, CT 06510, USA; Howard Hughes Medical Institute, School of Medicine, Stanford University, Stanford, CA 94305, USA

**Keywords:** Reciprocal synapse, dendro-dendritic synapse, neurexin (Nrxn), neuroligin (Nlgn), cerebellin (Cbln), compartmentalization

## Abstract

Reciprocal synapses are formed by neighboring dendritic processes that create the smallest possible neural circuit. Reciprocal synapses are widespread in brain and essential for information processing, but constitute a conceptual conundrum: How are adjacent pre- and post-synaptic specializations maintained as separate functional units? Here, we reveal an organizational principle for reciprocal synapses, using dendrodendritic synapses between mitral and granule cells in the mouse olfactory bulb as a paradigm. We show that mitral cells secrete cerebellin-1 to block the *cis*-interaction of mitral cell neurexins with neuroligins, thereby enabling their separate *trans*-interactions. Ablating either cerebellin-1 or neuroligins in mitral cells severely impaired granule cell→mitral cell synapses, as did overexpression of postsynaptic neurexins that form *cis*-complexes with neuroligins, but not of mutant neurexins unable to bind to neuroligins. Our data uncover a *cis*/*trans*-protein interaction network as a general design principle that organizes reciprocal dendro-dendritic synapses by compartmentalizing neurexin-based *trans*-synaptic protein complexes.

## INTRODUCTION

In connecting neurons into neural circuits, synapses exhibit a vast range of molecular and functional architectures. A prototypical synapse is formed by a presynaptic axonal bouton contacting a postsynaptic dendrite. Many synapses, however, do not conform to this design, for example double synapse spines (Knott et al., 2002), axo-axonic synapses (Ango et al., 2021; Cover & Mathur, 2021) and reciprocal dendro-dendritic synapses (Shepherd et al., 2020). Among these non-standard types of synapses, reciprocal dendro-dendritic synapses stand out because they comprise two antiparallel synaptic junctions formed by adjacent dendrites (Fig. 1A). The two dendrites generate both pre- and postsynaptic specializations, thereby assembling a two-neuron microcircuit, the smallest possible neural circuit that is broadly distributed throughout the brain.

**Figure 1.**
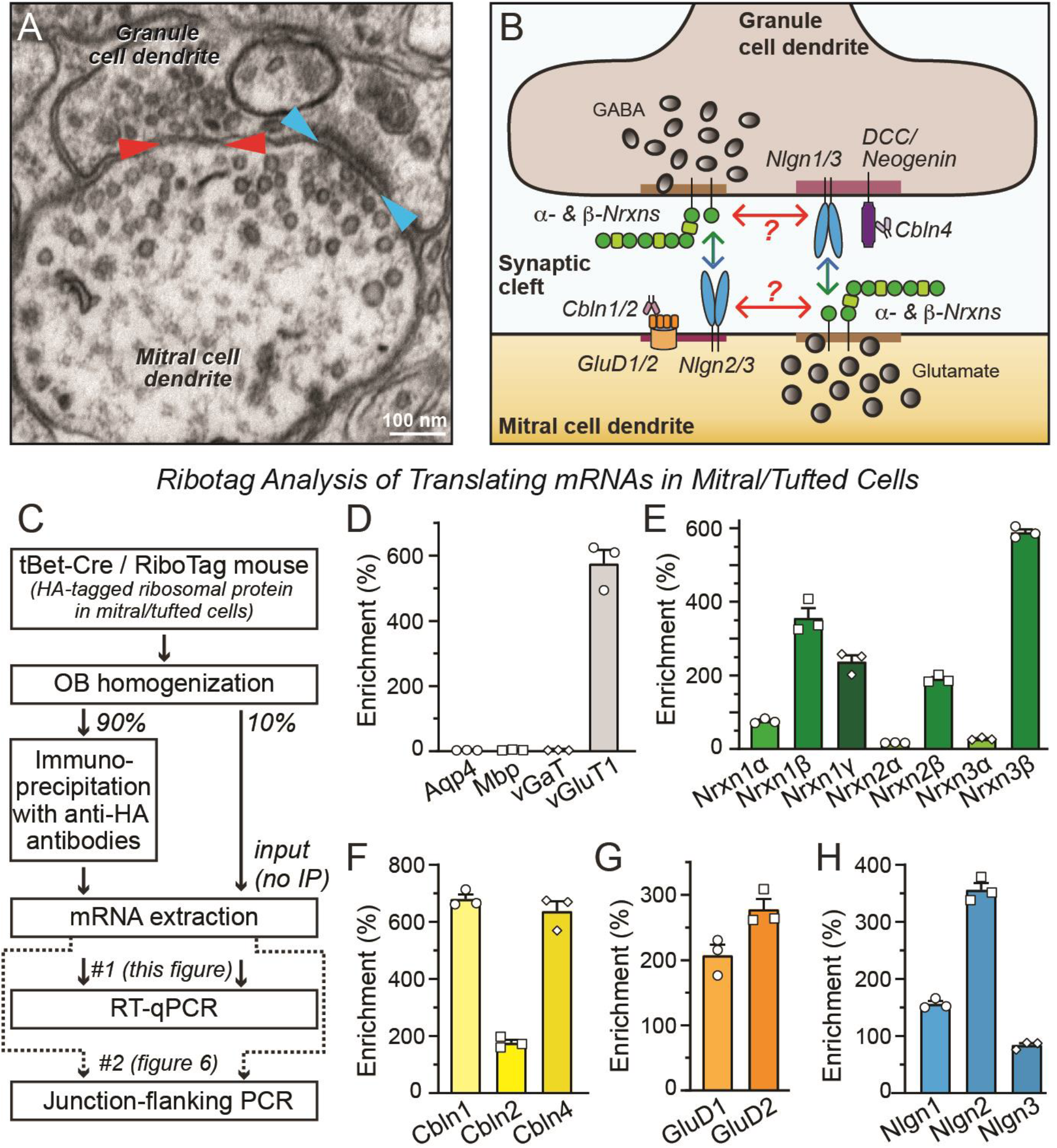
Mitral cells forming reciprocal dendro-dendritic synapses co-express neurexins and multiple neurexin ligands. ***A.*** Representative electron microscopy image of a reciprocal synapse in the olfactory bulb (red arrowhead, inhibitory granule cell→mitral cell (GC→MC) synapse; cyan arrowhead, excitatory mitral cell→granule cell synapse). ***B.*** Schematic of *trans*- and *cis*-interactions of neurexin-based synaptic adhesion molecules. Neurexins and their ligands are co-expressed in both mitral and granule cells. Due to the lack of physical compartmentalization, neurexins may interact with their ligands both in *cis* and *trans* configurations, with *cis*-interactions potentially inhibiting *trans*-interactions. The question of how two antiparallel neurexin signaling processes are organized to enable *trans*- over *cis*-interactions underlies the key to understand the design principle of reciprocal synapses. Note that in the diagram, the assignment of the pre- vs. postsynaptic localization of various neurexin ligands is for illustration purposes, and all ligands might actually be on both sides. ***C.*** Experimental strategy to isolated translating mRNAs from mitral and tufted cells using Ribotag mice and for analysis of these mRNAs. ***D.*** Summary graph demonstrating that mitral- and tufted-cell-specific mRNAs are de-enriched in mRNAs encoding aquaporin-4 (Aqp4, astrocyte marker), myelin basic protein (Mbp, oligodendrocyte marker, and vGaT (inhibitory neuron marker), but highly enriched in vGluT1 (excitatory neuron marker). ***E-H***. Summary graph demonstrating that mitral and tufted cells co-express specific isoform of neurexins (E), cerebellins (F), GluD1 and GluD2 (G), and neuroligins (H). All numerical data are means ± SEM (n=3 mice).

Reciprocal dendro-dendritic synapses were first observed between mitral/tufted cells and granule cells in the olfactory bulb (OB), where they constitute the most abundant type of synapse (Rall et al., 1966; reviewed in Urban & Arevian, 2009; Shepherd et al., 2020). OB reciprocal synapses are essential for olfactory information processing, and are subject to synaptic plasticity that mediates olfactory learning (Liu et al., 2017; Shepherd et al., 2020). Reciprocal dendro-dendritic synapses are also widely observed in other brain regions. They are found, among others, in the lateral geniculate nucleus (Famiglietti, 1970) the ventro-lateral nucleus of the thalamus (Harding, 1971), the motor cortex (Sloper and Powell, 1978), and the retina (Hartveit, 1999). Thus, reciprocal dendro-dendritic synapses represent constitutive building blocks of neural networks in the brain.

The structure of reciprocal dendro-dendritic synapses poses a unique cell-biological problem because each of the two participating dendrites forms pre- and post-synaptic specializations next to each other in the same plasma membrane domain (Fig. 1A). At a standard synapse, *trans-*synaptic signaling via adhesion molecules is thought to organize pre- and post-synaptic specializations. At reciprocal synapses, however, two antiparallel *trans-*synaptic signaling processes operate without physical compartmentalization. As a result, a presynaptic adhesion molecule in dendro-dendritic synapses, such as a neurexin, would have a high chance of interacting in *cis* with a postsynaptic ligand localized to the same membrane, such as neuroligins or cerebellins, instead of interacting with *trans*-synaptic ligands in the opposing membrane. To prevent such *cis*-dominant interactions, a mechanism must exist that compartmentalizes *trans*-interactions in favor of *cis*- interactions, but the nature of this mechanism is unknown. A similar need for compartmentalization exists for axo-axonic synapses that are also widespread in brain and contain adjacent pre- and postsynaptic specializations in the same plasma membrane.

Neurexins are among the best characterized presynaptic adhesion molecules for *trans-*synaptic signaling by interacting with multiple postsynaptic ligands, including neuroligins (Nlgns), cerebellins (Cblns, that in turn bind to GluD1/2 or to DCC/neogenin), LRRTMs, and dystroglycan (reviewed in Südhof, 2017; Kasem et al., 2018). In vertebrates, neurexins are encoded by three genes (*Nrxn1-3* in mice). Each neurexin gene expresses two principal forms, longer α-neurexins and shorter β-neurexins, that are transcribed from distinct promoters and are extensively alternatively spliced (Ushkaryov et al., 1992; Ushkaryov & Südhof, 1993; Tabuchi & Südhof, 2002). In addition, the *Nrxn1* gene has a third promoter for an even shorter isoform (Nrxn1γ ; Sterky et al., 2017). Neurexins are highly expressed in all neurons, but at different levels for each isoform. Whereas *trans*-synaptic signaling by presynaptic neurexins that engage postsynaptic ligands is conceptually straightforward in a prototypical synapse, in a reciprocal synapse such signaling poses a fundamental problem. Here, a presynaptic neurexin could potentially interact with its ligands in a *cis-* instead of a *trans-*configuration. The *cis*-interaction would be favored given the high local concentrations of neurexins and their ligands (Fig. 1B). Thus, the question arises: How are reciprocal synapses organized to prevent *cis*-interactions and favor *trans*-interactions of synaptic adhesion molecules? Insight into the molecular mechanisms that guide the functional compartmentalization of antiparallel neurexin signaling processes in reciprocal synapses will be crucial for progress in understanding the design principles of reciprocal synapses and other common but unconventional synapses.

To address this fundamental cell-biological question, we examined the function of neurexins and their ligands at reciprocal dendro-dendritic synapses in the OB. Surprisingly, neuroligins and Cbln1 are both required in mitral cells for granule cell→mitral cell (GC→MC) synaptic transmission, and both are essential for normal GABA receptor responses in mitral cells. We further identified that the mechanism behind the shared role of neuroligins and Cbln1 is that Cbln1 prevents mitral-cell neurexins from inhibiting the function of neuroligins in *cis,* allowing functional compartmentalization of two antiparallel neurexin signaling pathways in the reciprocal synapses.

## RESULTS

### Mitral cells of the OB co-express multiple neurexins and neurexin ligands

To determine whether neurexins and their neuroligin and cerebellin ligands are expressed by mitral/tufted cells that form dendro-dendritic synapses in the OB, we crossed Cre-dependent *RiboTag* mice with *tBet-Cre* mice (Sanz et al., 2009; Haddad et al., 2013). Using these mice, we selectively isolated mRNAs that were being translated in mitral/tufted cells, and performed quantitative RT-PCR on mitral/tufted cell-specific and total mRNA from the OB (Fig. 1C).

As expected, mRNAs encoding the excitatory neuron marker *vGluT1* were abundant in mitral/tufted cells, whereas mRNAs encoding the astrocyte marker *Agp4*, the oligodendrocyte marker *Mbp*, or the inhibitory neuron marker *vGaT* were largely absent (Fig. 1D). Consistent with previous *in situ* hybridization data (Ullrich, Ushkaryov & Südhof, 1995; Uchigashima et al., 2019), mitral/tufted cells expressed high levels of β-neurexins, with the highest enrichment observed for Nrxn3β (Fig. 1E). In contrast, α-neurexins were de-enriched (Fig. 1E). mRNAs encoding neuroligins, cerebellins, and the cerebellin receptors GluD1 and GluD2 were also abundant in mitral cells (Fig. 1F-1H). Cbln1 and Cbln4 mRNAs were highly enriched, whereas Cbln2 mRNAs were not as abundant (Fig. 1F). Cbln3 was not examined because of its exclusive expression in the cerebellum (Pang et al., 2000). Using single-molecule RNA *in-situ* hybridization, we confirmed the expression patterns of Cbln1, Cbln2 and Cbln4, and asked whether mitral and tufted cells equally synthesize various cerebellin isoforms. Strikingly, Cbln1 was more strongly expressed in mitral cells than in tufted cells, whereas Cbln4 exhibited the opposite pattern (Fig. 2A). Cbln2 was present at much lower levels in both mitral/tufted cells, but expressed at high levels in granule and periglomerular cells (Fig. 2A). Thus, mitral/tufted cells co-express multiple neurexins and neurexin ligands, suggesting that understanding the design principles of reciprocal synapses involves understanding the role of the neurexin-based protein interaction network at these synapses.

**Figure 2.**
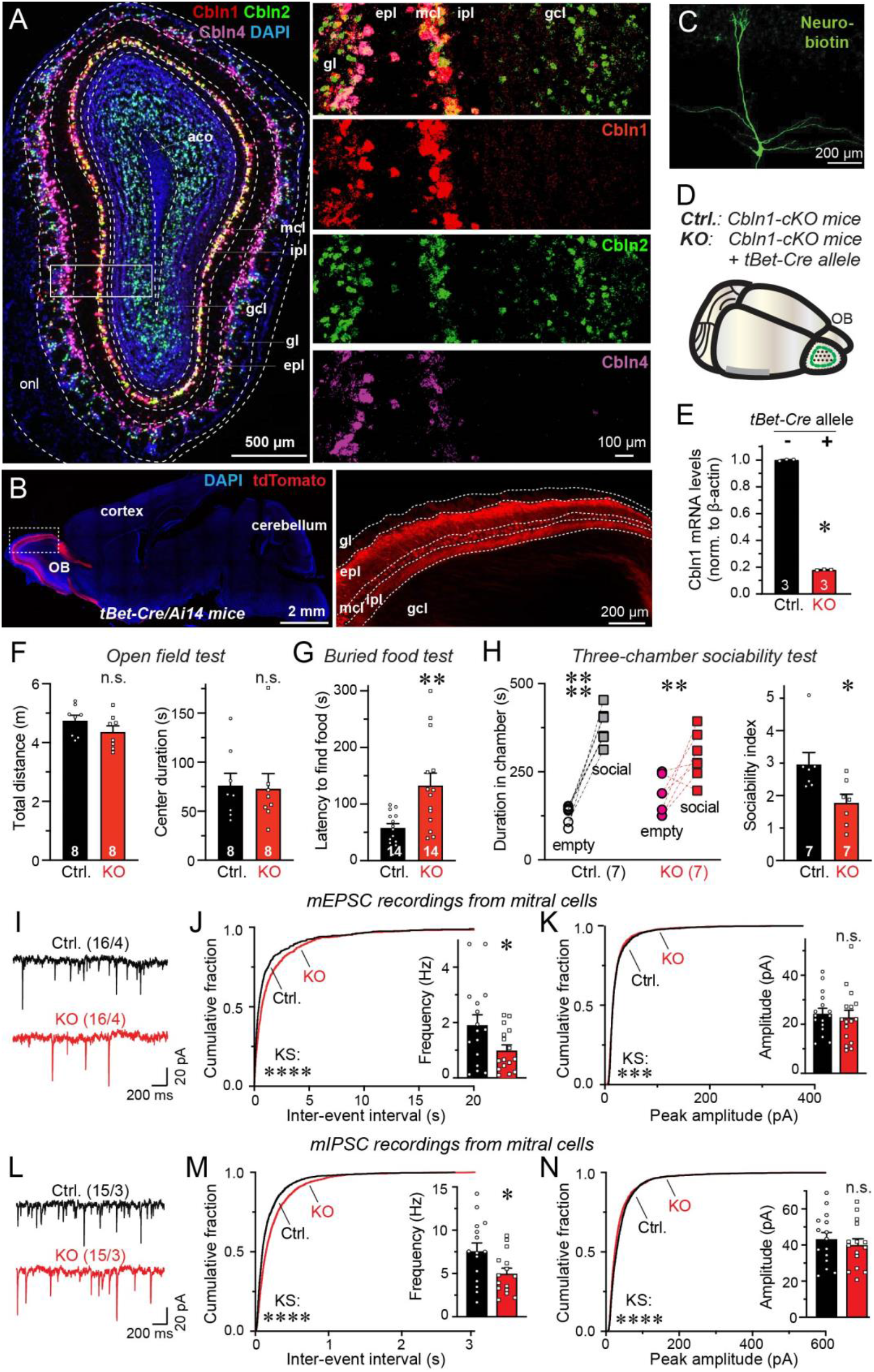
Deletion of *Cbln1* from mitral/tufted cells that predominantly express this isoform impairs olfactory behaviors and synaptic inputs onto mitral cells. ***A.*** Cbln1, Cbln2 and Cbln4 are differentially expressed in different cell types of the OB as revealed by single-molecule *in situ* hybridization (left, overview of an OB section stained for mRNAs encoding Cbln1 (red), Cbln2 (green) and Cbln4 (magenta) and for cell nuclei (DAPI, blue); right, higher-magnification views boxed in the left overview image, showing individual staining patterns). Abbreviations used: onl, olfactory sensory neuron layer; gl, glomerulus layer; epl, external plexiform layer; mcl: mitral cell layer; ipl: internal plexiform layer; gcl, granule cell layer; aco, anterior commissure of the olfactory limb. ***B.*** Validation of mitral- and tufted-cell-specific Cre expression in *tBet-Cre* mice that were crossed to *Ai14* indicator mice. Images depict Cre-dependent tdTomato expression in saggital sections of the OB (left, overview of the brain; right, higher magnification of the boxed OB area in the left image). ***C.*** Representative image of a mitral cell that was filled with neurobiotin via a patch pipette in an acute slice. ***D.*** Experimental strategy for the mitral and tufted cell-specific deletion of *Cbln1*. ***E.*** Cbln1 mRNA levels measured by quantitative RT-PCR in the OB of *Cbln1 cKO* mice that were crossed to *tBet-Cre* mice (normalized to actin mRNA). ***F-H***. Deletion of *Cbln1* from mitral and tufted cells selectively impairs olfactory behaviors (F, open field test; G, buried food test; H, three-chamber sociability test). **I**-***N***. Deletion of *Cbln1* decreases the frequency of miniature excitatory postsynaptic currents (mEPSCs; I-K) and of miniature inhibitory postsynaptic currents (mIPSCs; L-N) (I, L, representative traces; J, M, cumulative distributions of inter-event intervals [insets: bar graph of the mEPSC and mIPSC frequency]; K, N, cumulative distribution of mEPSC and mIPSC amplitudes [insets: bar graphs of the mEPSC and mIPSC amplitude]). All data are means ± SEM. Sample sizes are shown in the figures (E-H, number of mice) or representative traces (I & L, number of cells / number of mice). Statistical significance was assessed by Kolmogorov-Smirnov tests in the cumulative distributions of J-K and M-N, by *Student’s* t-test in the summary graphs of E-H, J-K and M-N, and by two-way ANOVA with Bonferroni’s multiple comparison test in the left panel of H (*, p<0.05; **, p<0.01; ***, p<0.001; ****, p<0.0001).

### Mitral-cell Cbln1 is essential for the organization of granule cell→mitral cell (GC→MC) synapses

Because of its robust expression, we first examined the role of Cbln1 in dendro-dendritic synapses of mitral cells. Mitral/tufted cells can be selectively manipulated in mice using Cre-recombinase expressed under the control of the *tBet* promoter (Haddad et al., 2013). Crosses of *tBet-Cre* mice with *Ai14* mice in which tdTomato expression is activated by Cre-mediated recombination (Madisen et al., 2010) validated exclusive expression of Cre in mitral/tufted cells of the OB (Fig. 2B). We filled patched mitral cells in acute OB slices with neurobiotin, which confirmed the typical cellular architecture of mitral cells (Fig. 2C). To generate mice with a mitral/tufted cell-specific deletion of *Cbln1*, we crossed *tBet-Cre* mice with *Cbln1* conditional KO (cKO) mice (Seigneur & Südhof, 2018) (Fig. 2D). The *Cbln1* deletion in mitral/tufted cells severely suppressed (∼90% decrease) the total Cbln1 mRNA levels in the OB, confirming that mitral/tufted cells express the vast majority of the Cbln1 in the OB (Fig. 2E). Behavioral experiments revealed that the mitral/tufted cell-specific deletion of *Cbln1* significantly impaired a mouse’s foraging behavior as assayed by the buried food test (Fig. 2G). The mitral/tufted cell deletion of *Cbln1* also decreased the sociability of mice without affecting their general mobility or increasing their anxiety levels (Fig. 2F-H). Thus, mitral/tufted cell expression of Cbln1 is essential for olfactory behaviors.

We next examined the effect of the mitral cell *Cbln1* deletion on synaptic transmission. Whole-cell patch-clamp recordings in acute slices revealed that the *Cbln1* deletion reduced the frequency of both mIPSCs (∼35%) and mEPSCs (∼50%) without significantly altering their amplitudes or kinetics (Fig. 2I-N; S1A-D). The *Cbln1* deletion did not affect the capacitance or input resistance of mitral cells, suggesting that the cells were healthy (Fig. S1E-F). Thus, postsynaptic Cbln1 is required for organizing both inhibitory and excitatory inputs onto mitral cells.

The postsynaptic requirement of Cbln1 for the function of mitral cell synapses is surprising because Cbln1 is thought to regulate excitatory and inhibitory synapse formation by a presynaptic mechanism (Matsuda et al., 2010; Uemura et al., 2010; Ito-Ishida et al., 2012 and 2014; Ibata et al., 2019), although we recently detected a postsynaptic action of Cbln2 in excitatory hippocampal synapses (Dai et al., 2021). Since most mIPSCs in mitral cells are derived from dendro-dendritic GC→MC synapses, the decrease in mIPSC frequency suggests that Cbln1 functions in GC→MC synapses. To test this notion, we recorded evoked IPSCs in mitral cells that were induced by stimulating the apical dendrites of granule cells (Fig. 3A). We used input/output measurements to control for the variability in the placement of the stimulating electrode. These experiments revealed that the *Cbln1* deletion caused a major decrease (∼45%) in GC→MC synaptic strength (Fig. 3B-D). In addition, the *Cbln1* deletion increased the rise but not decay times of GC→MC IPSCs (Fig. 3E-F).

**Figure 3.**
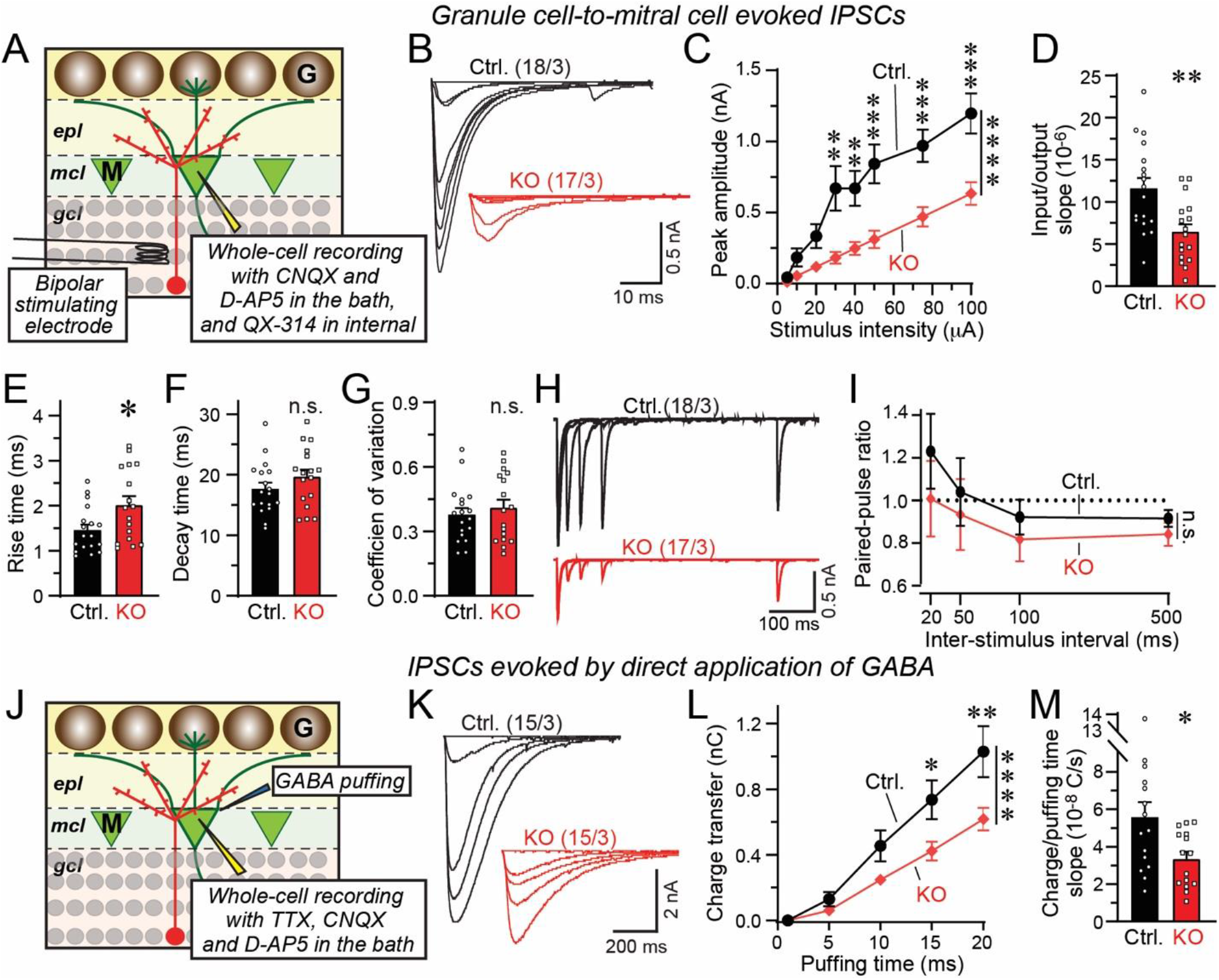
Mitral-cell Cbln1 enables GC→MC synaptic transmission by a postsynaptic mechanism regulating GABA_A_-receptors. ***A***. Experimental paradigm for recordings of IPSCs evoked by stimulation of granule cell dendrites using an extracellular bipolar electrode. ***B-D***. The *Cbln1* deletion in mitral and tufted cells severely impairs GC→MC synaptic transmission. Recordings are from acute OB slice from *Cbln1 cKO* mice crossed with *tBet-Cre/+; Cbln1-cKO* mice (B, representative traces; C, input/output curve of GC→MC IPSC peak amplitudes as a function of the stimulus intensity; D, summary of the slope of the input/output curves recorded in individual experiments). ***E & F***. The *Cbln1* deletion slows down the kinetics of GC→MC IPSCs in mitral cells (E, rise time; F, decay time; both measured at a 75 µA stimulus intensity). ***G***. The *Cbln1* deletion has no effect on the coefficient of variation (G) or the paired-pulse ratio (H & I) of GC→MC IPSCs in mitral cells (G, summary graph of the coefficient of variation; H & I, representative traces and summary plot of the paired pulse ratio; all monitored at a 75 µA stimulus intensity). ***J***. Experimental paradigm for recordings of IPSCs evoked by direct application of GABA to mitral cells using a Picospritzer. ***K-M***. The *Cbln1* deletion severely impairs the IPSCs elicited in mitral cells by direct application of GABA (K, representative traces; L, summary plot of the total IPSC charge elicited by different application durations of GABA (10 mM); M, summary graph of the slope of the GABA-response curve recorded in individual experiments). All numerical data are means ± SEM (numbers of cells/mice analyzed are indicated above the sample traces). Statistical significance was assessed by *Student’s* t-test in D-G and M, and by two-way ANOVA with Bonferroni’s multiple comparison test in C, I and L (*, p<0.05; **, p<0.01; ***, p<0.001; ****, p<0.0001).

Most inhibitory inputs onto mitral cells are provided by granule cells, but periglomerular neurons, short-axon cells, and other types of interneurons also form inhibitory synapses on mitral cells (Eyre et al., 2008; Zhou et al., 2020). To reveal whether the phenotype of the *Cbln1* deletion at inhibitory synapses is specific for GC→MC synapses or whether it broadly applies to all inhibitory synapses of mitral cells, we measured IPSCs evoked by stimulating inhibitory periglomerular inputs (Fig. S2). We detected only a small increase in synaptic strength that was not statistically significant when the slopes of input/output curves were calculated (Fig. S2C-D). Thus, the *Cbln1* deletion does not uniformly impair all inhibitory synapses of mitral cells.

### Deletion of *Cbln1* decreases GABA-induced IPSCs but not GABA release

At least three mechanisms could account for the synaptic impairment induced by the *Cbln1* deletion: A decrease in release probability at GC→MC synapses, a decline in postsynaptic receptor responses, or a reduction in synapse numbers. To distinguish between these three hypotheses, we examined the coefficient of variation (CV) and the paired-pulse ratio (PPR) of evoked IPSCs as indirect but sensitive measures of release probability (both monitored at a 75 µA stimulus intensity). The *Cbln1* deletion had no effect on either parameter in GC→MC synapses (Fig. 3G-I), suggesting that the release probability was unchanged.

Next, we explored the possibility that the *Cbln1* deletion decreases postsynaptic GABA_A_-receptor responses at GC→MC synapses. For this purpose, we directly ‘puffed’ GABA onto the mitral cell soma using a Picospritzer, and monitored the resulting IPSCs (Fig. 3J). Strikingly, we observed a robust reduction (∼40%) in GABA-induced IPSCs in *Cbln1*-deficient mitral cells (Fig. 3K-M). This phenotype is consistent with the notion that the *Cbln1* deletion suppresses postsynaptic GABA_A_-receptor responses, but does not rule out a decrease in inhibitory synapse numbers, which would also decrease the overall GABA_A_-receptor response. Moreover, the observed decrease in mIPSC frequency in *Cbln1*-deficient mitral cells (Fig. 2M) would also be consistent with either a lowered GABA_A_-receptor responsiveness or a decrease in synapse numbers because a decreased GABA response would enhance ‘silent’ inhibitory synapses and make mIPSCs less detectable. Thus, we needed to determine whether the *Cbln1* deletion induced a synapse loss in the OB, as had been observed for the *Cbln1* deletion in the cerebellum (Hirai et al., 2005; Uemura et al., 2010).

### The *Cbln1* deletion has no effect on synapse numbers or ultrastructure

Using cryosections from mice with a *Cbln1* deletion in mitral/tufted cells, we stained dendro-dendritic synapses on mitral cells for vGaT as a universal inhibitory synapse marker and for synaptophysin-2 as a specific GC→MC synapse marker in the OB (Bergmann et al., 1993) (Fig. 4A-B). We observed no change in the number or size of inhibitory synapses in the external plexiform layer of mice with a *Cbln1* deletion in mitral/tufted cells, suggesting that the *Cbln1* deletion does not alter inhibitory synapse numbers (Fig. 4A-F).

**Figure 4.**
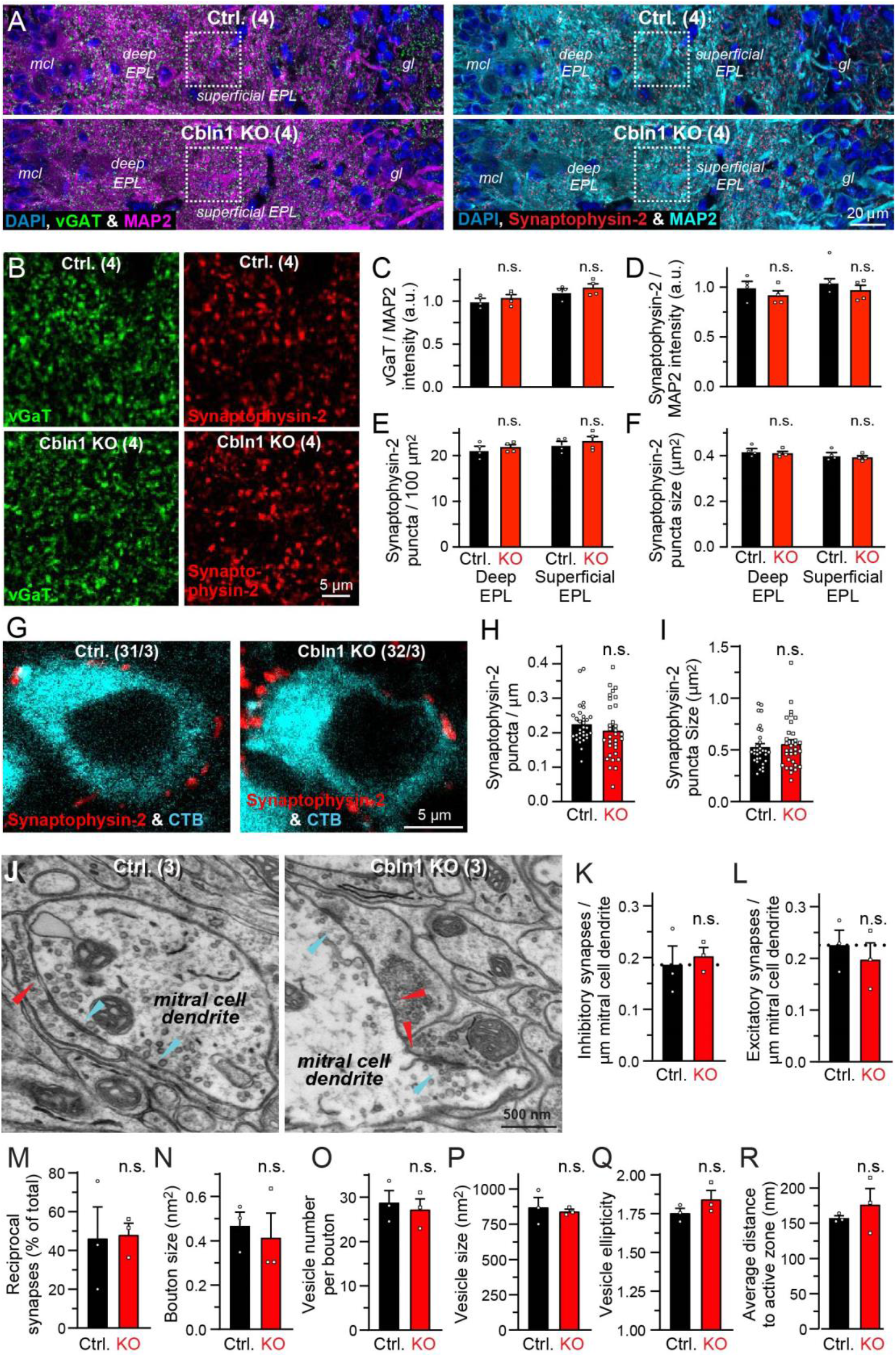
Deletion of *Cbln1* from mitral and tufted cells does not change the density or ultrastructure of dendro-dendritic synapses. ***A-F***. The *Cbln1* deletion from mitral cells does not decrease the density of vGAT- or synaptophysin-2-positive synapses on mitral cells (A & B, representative low and higher magnification images of of OB sections stained for vGaT, Synaptophysin-2 and MAP2 as indicated; C-F, summary graphs of the intensity of vGAT staining (C) and the density, staining intensity, and size of synaptophysin-2-positive puncta (D-F)). Note that synaptophysin-2 is a relatively specific marker for presynaptic dendro-dendritic specializations in the OB. Abbreviations: epl, external plexiform layer; mcl, mitral cell layer; gl, glomerular layer. ***G-I***. Immunocytochemistry for perisomatic synaptophysin-2 in retrogradely labeled mitral cells without or with the *Cbln1* KO document that the *Cbln1* KO does not decrease the number of inhibitory perisomatic synapses onto mitral cells. Mitral cells were labeled by injection of fluorescent chloral toxin B (CTB) into the piriform cortex (G, presentative images; H & I, summary graphs of the density and size of synaptophysin-2-positive puncta). ***J-R***. Analysis of reciprocal dendro-dendritic synapses in the external plexiform layer of the OB by electron microscopy shows that the *Cbln1* KO does not decrease the density or change the ultrastructure of dendro-dendritic synapses (J, representative images [red arrows, symmetric GC→MC) synapses; cyan arrow, asymmetric mitral cell to granule cell synapses]; K & L, summary graphs of the GC→MC and the mitral cell→granule cell synapse densities normalized for the mitral cell dendrite length; M, summary graph of the percentage of reciprocal synapses among GC→MC synapses; N-R, summary graphs of the bouton size (N), vesicle number per bouton (O), vesicle size (P), vesicle ellipticity (Q), and distance of the vesicles to the active zone (R) of symmetric GC→MC synapses. All numerical data are means ± SEM. Sample sizes are shown in the representative images as in the number of mice in A-B & J and the number of cells from three mice in G. Statistical analyses were performed by *Student’s* t-test in H-R, and by two-way ANOVA with Bonferroni’s multiple comparison test in C-F (n.s.: p>0.05).

Some granule cells form synapses on the soma of mitral cells (Naritsuka et al., 2009) that is also targeted in our GABA puffing experiment. Therefore, a specific loss of somatic inhibitory synapses could be caused by the *Cbln1* deletion. To test this possibility, we specifically analyzed perisomatic synapses of mitral cells. We retrogradely labeled the mitral cell soma with fluorescently tagged cholera toxin B (CTB) that was injected into the piriform cortex, and stained presynaptic granule cell inputs using synaptophysin-2 staining. However, we again observed no change in synapse numbers on *Cbln1*-deficient mitral cells (Fig. 4G-I).

Finally, to independently confirm the immunocytochemical conclusions and to test whether the *Cbln1* deletion affects the structure of dendro-dendritic synapses, we analyzed them by electron microscopy (EM) (Fig. 4J). The *Cbln1* deletion in mitral cells produced no change in the density of dendro-dendritic synapses on mitral cell dendrites, confirming the immunocytochemical results (Fig. 4K-M). Moreover, we detected no change in any structural parameter of synapses, such as the size of presynaptic terminals (Fig. 4N), the number of vesicles per terminal (Fig. 4O), the size of vesicles (Fig. 4P), the ellipticity of vesicles (Fig. 4Q), or the average distance of vesicles to the active zone (Fig. 4R).

Thus, Cbln1 is essential in mitral cells as a postsynaptic regulator of GC→MC synapses that controls the GABA_A_-receptor responses of mitral cells without affecting either the release probability or number of GC→MC synapses.

### Neuroligins are also essential for GC→MC synaptic transmission

Since both neuroligins and Cbln1 are ligands for neurexins and since neuroligins are also abundantly expressed in mitral cells (Fig. 1H; Varoqueaux et al., 2004), we next asked whether neuroligins contribute to the function of GC→MC synapses in the OB. In mammals, neuroligins are encoded by four genes, *Nlgn1-4* (Ichtchenko et al., 1995 and 1996; Nguyen & Südhof, 1997; Bolliger et al., 2008). To test whether neuroligins function at GC→MC synapses, we utilized *Nlgn1234* quadruple conditional KO (qcKO) mice. In these mice, the *Nlgn1*, *Nlgn2*, and *Nlgn3* genes are floxed and can be inactivated by Cre recombinase, whereas the *Ngln4* gene is constitutively deleted (Zhang et al., 2015, 2016 and 2018; Chanda et al., 2017). We utilized these mice for the analysis of neuroligin function in mitral cells instead of individual neuroligin KO mice because multiple neuroligins often operate in the same synapse and are partly redundant (Chanda et al., 2017), and because analysis of individual KO mice would be prohibitively time-consuming.

Using stereotactic injections, we introduced into the piriform cortex of *Nlgn1234* qcKO mice a retrogradely transported adeno-associated virus (rAAV2-retro; Tervo et al., 2016) that co-expresses EGFP with Cre or ΔCre (mutant inactive Cre as a control) (Fig. 5A). rAAV2-retro infects axons in the piriform cortex that emanate from mitral/tufted cells in the OB, thereby enabling selective expression of Cre or ΔCre in mitral/tufted cells of the OB. Two to three weeks after injections, we analyzed the infected mitral cells in the OB electrophysiologically (Fig. 5B).

**Figure 5.**
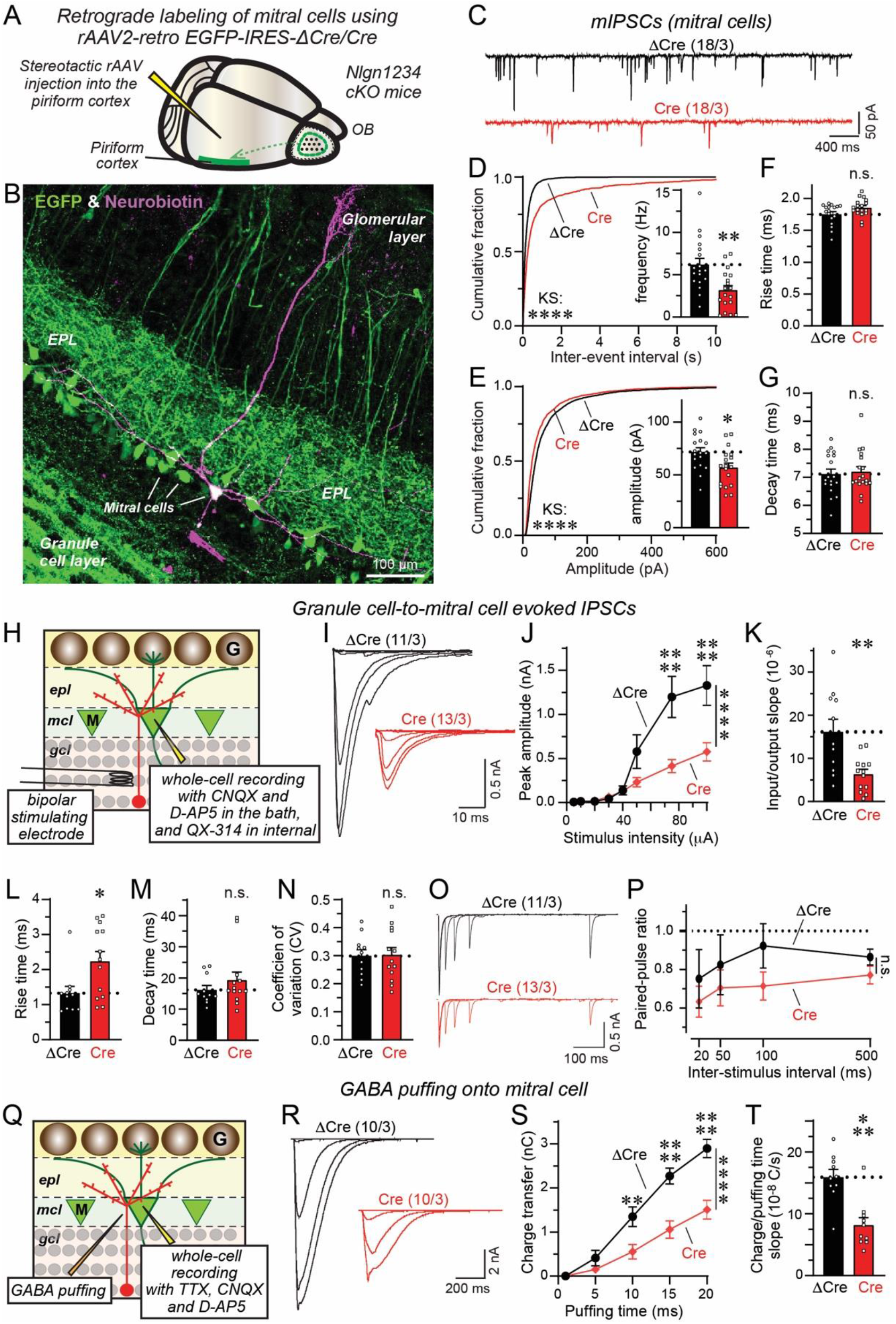
Deletion of all neuroligins severely impairs GC→MC synaptic transmission by suppressing the postsynaptic GABA-responsiveness of mitral cells similar to the *Cbln1* deletion. ***A.*** Experimental strategy for selective expression of Cre (or ΔCre as a control) in mitral cells of the OB by injecting rAAV2-retro co-expressing EGFP with Cre or ΔCre into the piriform cortex. rAAV2-retro is taken up by mitral cell axons projecting to the piriform cortex, resulting in selective infection of mitral cells in the OB. ***B.*** Representative image of an OB section with mitral cells expressing EGFP (green) via retrograde infection with rAAV2-retro in the piriform cortex. A mitral cell was patched and filled with neurobiotin (magenta). Abbreviation: epl, external plexiform layer. ***C-G***. The deletion of all neuroligins from mitral cells using rAAV2-retro infections in conditional quadruple *Nlgn1234* KO mice decreases the mIPSC frequency and amplitude (C, representative mIPSC traces; D, cumulative distribution of mIPSC inter-event intervals and summary graph of the mIPSC frequency; E, cumulative distribution of mIPSC amplitudes and summary graph of the mIPSC amplitude; F & G, summary graphs of the mIPSC rise (10% to 90%) and decay times (90% to 10%). ***H***. Experimental paradigm for recordings of IPSCs evoked by stimulation of granule cell dendrites using an extracellular bipolar electrode. ***I-K***. The deletion of all neuroligins severely impairs GC→MC synaptic transmission (I, representative traces; J, input/output curve of GC→MC IPSC peak amplitudes as a function of the stimulus intensity; K, summary of the slope of the input/output curves recorded in individual experiments). ***L & M***. The neuroligin deletion slows down the kinetics of GC→MC IPSCs in mitral cells (L, rise time; M, decay time; both measured at a 75 µA stimulus intensity). ***N-P***. The neuroligin deletion has no effect on the coefficient of variation (N) or the paired-pulse ratio (O & P) of GC→MC IPSCs in mitral cells (N, summary graph of the coefficient of variation; O & P, representative traces and summary plot of the paired pulse ratio; all monitored at a 75 µA stimulus intensity). ***Q***. Experimental paradigm for recordings of IPSCs evoked by direct application of GABA to mitral cells using a Picospritzer. ***R-T***. The deletion of all neuroligins from mitral cells greatly decreases the magnitude of IPSCs elicited by direct application of GABA (R, representative traces; S, summary plot of the total IPSC charge elicited by different application durations of GABA (10 mM); T, summary graph of the slope of the GABA-response curve recorded in individual experiments). All numerical data are means ± SEM (numbers of cells/mice analyzed are indicated above the sample traces). Statistical analyses were performed by Kolmogorov-Smirnov tests in the cumulative distributions of D-E, by *Student’s* t-test in the summary graphs of D-G, K-N and T, and by two-way ANOVA with Bonferroni’s multiple comparison test in J, P and S (*, p<0.05; **, p<0.01; ***, p<0.001; ****, p<0.0001).

Recordings of mIPSCs from mitral cells showed that the deletion of all neuroligins produced a large reduction (∼50%) in mIPSC frequency and a significant decrease (∼20%) in mIPSC amplitude without changing the mIPSC kinetics (Fig. 5C-G). As discussed above, most mIPSCs in mitral cells are derived from dendro-dendritic GC→MC synapses, suggesting that the neuroligin deletion impairs these synapses. Indeed, analysis of evoked GC→MC IPSCs revealed that the pan-neuroligin deletion caused a large reduction (∼60%) in GC→MC IPSC amplitudes and a significant increase in IPSC rise times but not decay times (Fig. 5H-M). At the same time, the neuroligin deletions had no effect on the membrane capacitance and input resistance of mitral cells, suggesting that it did not alter their size or viability (Fig. S3).

We next asked whether the neuroligin deletion reduces GC→MC synaptic transmission by a mechanism similar to that of the *Cbln1* deletion. The neuroligin deletion had no effect on the coefficient of variation (CV) and paired-pulse ratio (PPR) of evoked IPSCs (Fig. 5N-P), suggesting that the release probability was unchanged. Similar to the *Cbln1* deletion, however, the neuroligin deletion robustly reduced (∼50%) the amplitude of IPSCs evoked by direct application of GABA onto the mitral cell soma using a picospritzer (Fig. 5Q-T). Thus, the pan-neuroligin deletion suppresses postsynaptic GABA_A_-receptor responses without affecting the presynaptic release probability, suggesting that neuroligins are essential organizers of GC→MC synapses in the OB similar to Cbln1.

### Alternative splicing of postsynaptic neurexins at SS4 in mitral cells regulates GC→MC synapses

Why are *Cbln1* and neuroligins both required for organizing GC→MC synapses, even though they bind to the same domain of neurexins? This requirement could operate by two mechanisms: A parallel involvement of *Cbln1* and neuroligin interacting with granule-cell neurexins in GC→MC synapses, or a non-canonical function of Cbln1 or neuroligins that indirectly renders GC→MC synapses functional. This question is of general interest given that most neurons co-express cerebellins and neuroligins, but is particularly important for the reciprocal synapses in which neurexins and neurexin ligands can interact in both *cis* and *trans* configurations (Fig. 1B).

If mitral cell Cbln1 acted directly in GC→MC synapses, it would have to function by binding to presynaptic granule cell neurexins, whereas a non-canonical postsynaptic function would involve a binding to postsynaptic neurexins. Interestingly, measurements of neurexin alternative splicing in mitral cell mRNAs using junction-flanking RT-PCR showed that all neurexins are predominantly expressed in mitral/tufted cells as splice site 4 containing (SS4+) variants (Fig. 6A-B). This is important because only SS4+ variants of neurexins can bind Cbln1, whereas SS4− variants cannot (Uemura et al., 2010; Joo et al., 2011; Matsuda & Yuzaki, 2011).

**Figure 6.**
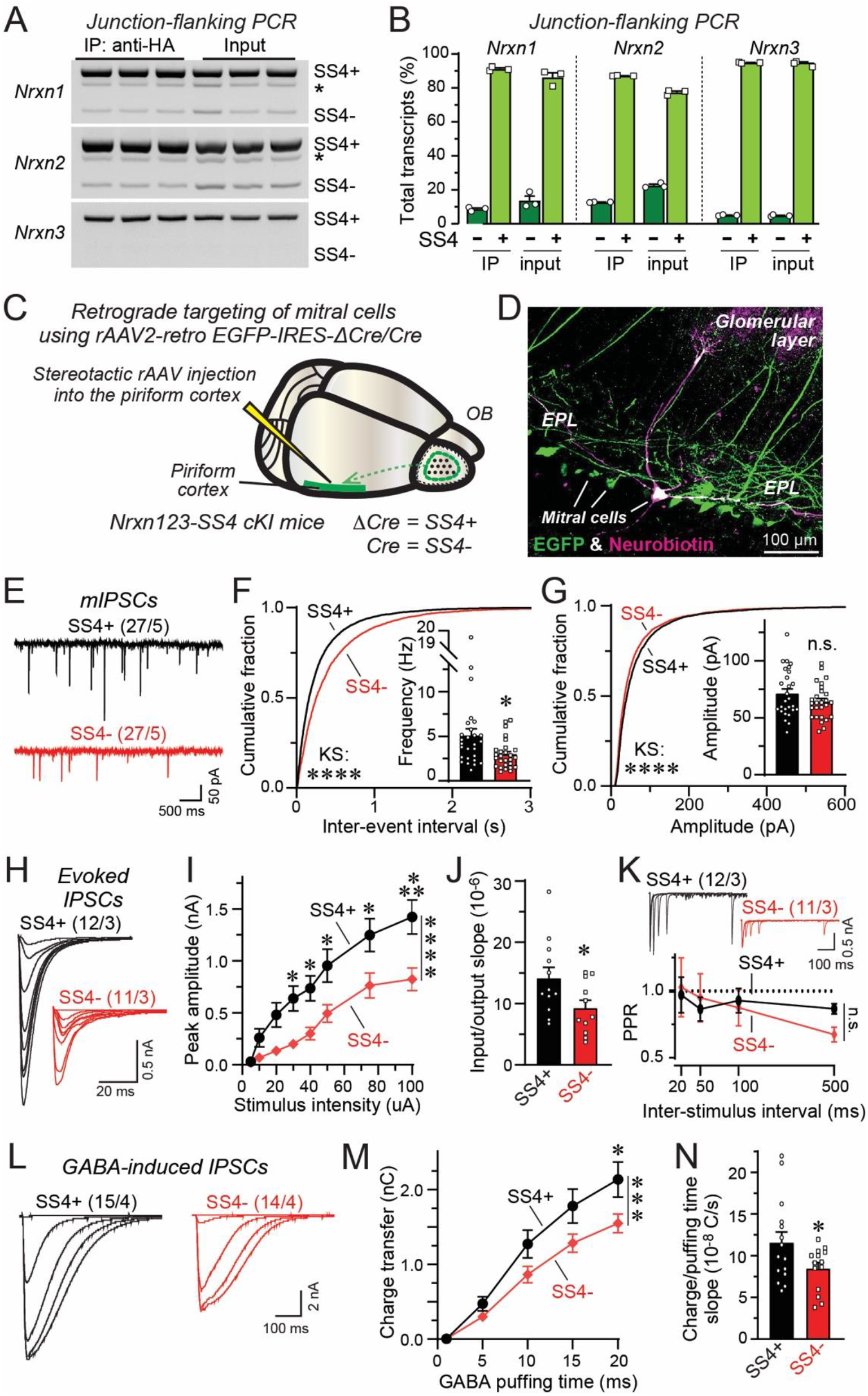
Converting mitral cell SS4+ neurexins into SS4− neurexins severely impairs GC→MC synaptic transmission by a mechanism recapitulating that of the *Cbln1* and *Nlgn1234* deletions. ***A & B***. Nrxn1, Nrxn2, and Nrxn3 are predominantly expressed as SS4+ variants in mitral/tufted cells of the OB (A, representative image of junction-flanking PCR analyses of total OB mRNA [input] and mitral/tufted cell mRNA isolated using RiboTag pulldowns [IP: anti-HA; see Figure 1C]; * represents unspecific band determined by band size; B, summary graph of the SS4 splicing profile of mitral/tufted-cell [IP: anti-HA] and whole-OB translating mRNA [input]). ***C.*** Experimental strategy for controlling SS4 alternative splicing of neurexins in in mitral cells. Triple conditional knockin (cKI) mice in which all three neurexins are constitutively expressed as SS4+ variants but can be converted into SS4− variants (Dai et al., 2019) are injected into the piriform cortex with rAAV2-retro co-expressing EGFP with Cre (to convert the SS4+ neurexins into SS4− neurexins) or ΔCre (as a control). Mitral cells are analyzed 2-3 weeks later in acute slices by whole-cell patch-clamp recordings. ***D.*** Representative image of a recorded mitral cell with filled neurobiotin and EGFP expressed by retrogradely transported rAAV2-retro. ***E-G***. Postsynaptic conversion of SS4+ into SS4− neurexins in mitral cells decreases the mIPSC frequeny (E, representative mIPSC traces; F, cumulative distribution of mIPSC inter-event intervals and summary graph of the mIPSC frequency; G, cumulative distribution of the mIPSC amplitudes and summary graph of the mIPSC amplitude). ***H-K***. Postsynaptic conversion of SS4+ into SS4− neurexins in mitral cells suppresses the amplitude of evoked GC→MC IPSCs without changing their paired-pulse ratio (H, representative traces of GC→MC IPSCs; I & J, input/output curve analyzing the GC→MC IPSC peak amplitude as a function of stimulus intensity (I), and summary graph of the input/output slopes (J); K, representative traces (top) and summary plot (bottom) of the IPSC paired-pulse ratio monitored at a 75 µA stimulus intensity). ***L-N***. Postsynaptic conversion of SS4+ into SS4− neurexins reduces the response of mitral cells to directly applied GABA (L, representative traces of IPSCs induced by direct GABA applications; M, input/output curve measuring the GABA-induced charge transfer as a function of the duration of the GABA application; N, summary graph of the slope of the input/output curves). All numerical data are means ± SEM (numbers of cells/mice analyzed are indicated above the sample traces in E-N). N=3 mice were analyzed in A-B. Statistical analyses were performed by Kolmogorov-Smirnov tests in the cumulative distributions of F-G, by *Student’s* t-test in the summary graphs of F-G, J and N, and by two-way ANOVA with Bonferroni’s multiple comparison test in I, K and M (*, p<0.05; **, p<0.01; ***, p<0.001; ****, p<0.0001).

Motivated by this finding, we tested whether converting the SS4+ variants of neurexins in mitral cells into SS4− variants that blocks their binding to Cbln1 but not to neuroligins suppresses GC→MC synaptic transmission similar to the *Cbln1* deletion. For this purpose, we used triple-conditionally mutant mice that constitutively express all neurexins as SS4+ variants (*Nrxn123-SS4+* cKI mice; Dai et al., 2019). In these mice, Cre recombinase converts the SS4+ neurexin variants into SS4− variants. We injected rAAV2-retro’s co-expressing EGFP with ΔCre (control, retains SS4+ neurexin expression) or Cre (converts SS4+ neurons into SS4− neurexins) into the piriform cortex of *Nrxn123-SS4+* cKI mice, and analyzed EGFP-positive mitral cells in acute slices by whole-cell patch-clamp recordings two weeks later (Fig. 6C-D).

The conversion of postsynaptic SS4+ to SS4− neurexins in mitral cells robustly decreased the mIPSC frequency (∼40%) without inducing major changes in mIPSC amplitude or mIPSC kinetics (Fig. 6E-G, S4A-B). Moreover, the SS4 conversion produced a highly significant reduction (∼35%) in the amplitude of evoked GC→MC IPSCs, and increased both the rise and decay times of evoked IPSCs (Fig. 6H-J; S4C-D). No change in the coefficient of variation or paired-pulse ratio was observed (Fig. 6K; Fig. S4E). Similar to the deletion of *Cbln1* or neuroligins in mitral cells, the conversion of SS4+ to SS4− neurexins in mitral cells also decreased the amplitude of IPSCs induced by direct application of GABA (Fig. 6L-N; ∼25% decrease). In none of these experiments, a change in the mitral cell membrane properties was observed (Fig. S4F-G).

The phenotype of the conversion of SS4+ to SS4− neurexins in mitral cells indicates that, surprisingly, Cbln1-binding to neurexins acts postsynaptically in dendro-dendritic GC→MC synapses. However, the results do not completely rule out the possibility that Cbln1 also acts by binding to presynaptic neurexins in GC→MC synapse, i.e. that the Cbln1/neurexin and neuroligin/neurexin signaling pathways operate in the same synapse simultaneously. To test this possibility, we converted SS4+ neurexins in granule cells into SS4− neurexins. We expressed Cre or ΔCre (as a control) in the OB of *Nrxn123*-SS4+ cKI mice using AAV_DJ_‘s that under the conditions used only infect granule cells in the OB (Fig. S4H-I). Rendering all granule cell neurexins SS4− had no effect on evoked GC→MC IPSCs, indicating that presynaptic neurexins in GC→MC synapses do not functionally require Cbln1 binding (Fig. S4J-S). Moreover, CRISPR-mediated deletion of postsynaptic GluD1 in mitral cells had no effect on GC→MC synapses (Fig. S5). Admittedly, this lack of an effect might have been due to redundancy between GluD1 and GluD2 because of GluD2 enrichment in mitral/tufted cells (Fig. 1F). However, it was shown that GluD1 rather than GluD2 preferentially induces inhibitory presynaptic differentiation (Yasumura et al., 2012); and that GluD2 deletion increases rather than decreases inhibitory synapse number in the cerebellum (Ito-Ishida et al., 2014). Viewed together, these observations argue against the possibility that Cbln1 mediates GC→MC synapses through a transsynaptic complex involving granule-cell neurexins and mitral-cell GluD1.

### Neurexin/neuroligin *cis*-interactions inhibit neuroligin *trans*-interactions in a manner that is alleviated by *Cbln1*

Viewed together, our data suggest that Cbln1 and neuroligin function postsynaptically in GC→MC synapses, and that this function requires postsynaptic SS4+ variants of neurexins and is abolished by SS4− variants of neurexins. A plausible model to explain these data is that Cbln1 is essential in dendro-dendritic GC→MC synapses because it blocks a *cis*-neurexin/neuroligin interaction that would otherwise inhibit postsynaptic neuroligins in this synapse. This model implies that Cbln1 competitively blocks neuroligins from binding to neurexins. To test this implication, we compared binding of Cbln1 tagged with an HA epitope (HA-Cbln1) or of the extracellular domains of Nlgn1 fused to the IgG Fc-domain (Nlgn1-ECD-Fc) to the two SS4 splice variants of Nrxn1β (Nrxn1β^SS4+^ and Nrxn1β^SS4−^) (Fig. 7A). As expected, HA-Cbln1 only bound to Nrxn1β^SS4+^ but not to Nrxn1β^SS4−^, whereas Nlgn1-ECD-Fc bound to both (Fig. 7B). When HA-Cbln1 and Nlgn1-ECD-Fc were added sequentially, HA-Cbln1 blocked binding of Nlgn1-ECD-Fc to Nrxn1β^SS4+^, independent of whether Cbln1 was bound to Nrxn1β^SS4+^ first, or added after Nlgn1-ECD-Fc had been prebound (Fig. 7B). HA-Cbln1, however, had no effect on Nlgn1-ECD-Fc binding to Nrxn1β^SS4−^ that is unable to bind cerebellins (Fig. 7B). Consistent with previous binding affinity measurement (Araç et al., 2007; Chen et al., 2008; Joo et al., 2011), these results show that Cbln1 binds more tightly to Nrxn1β^SS4+^ than Nlgn1.

**Figure 7.**
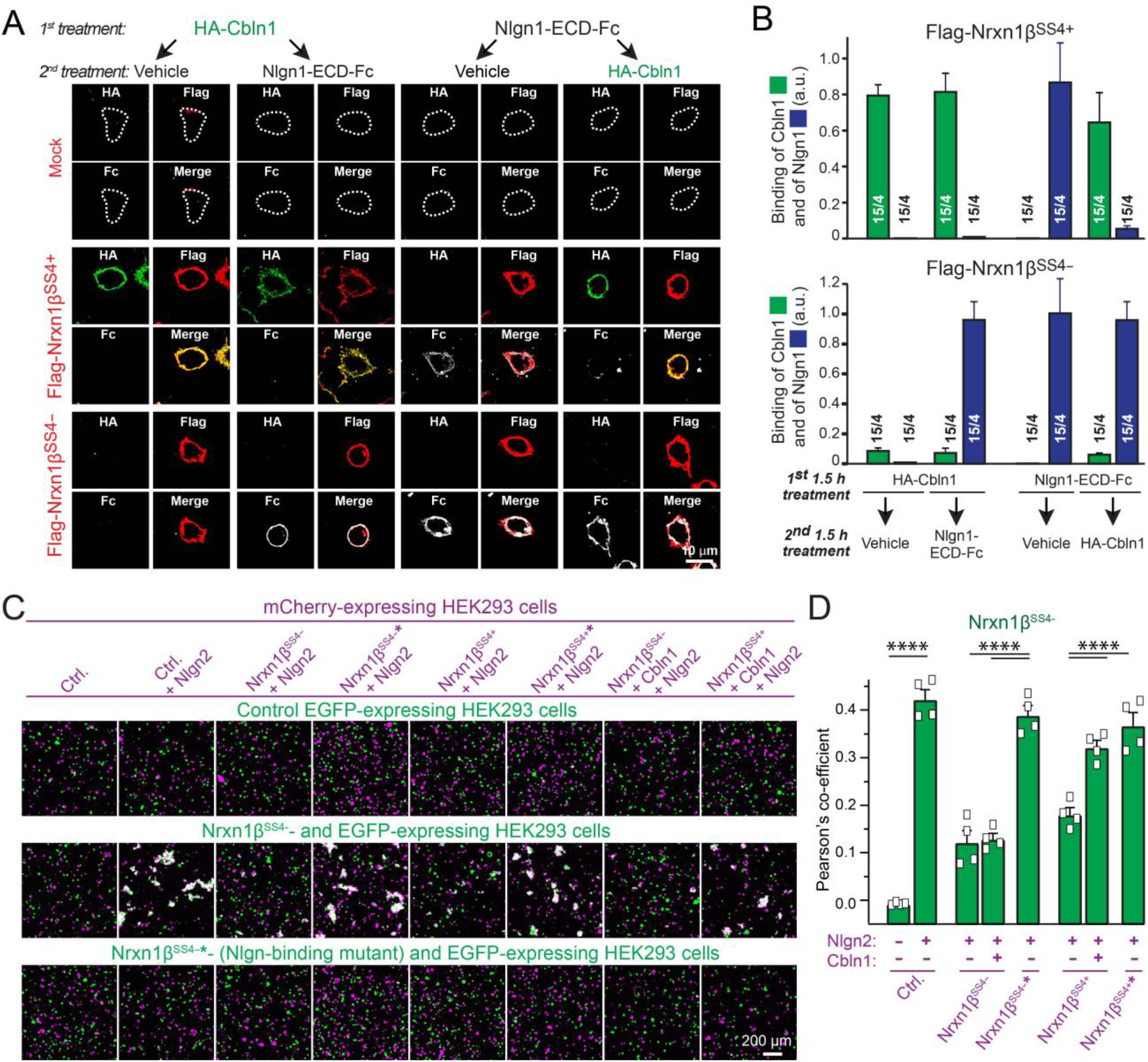
Cbln1 displaces Nlgn1 from the SS4+ but not the SS4− splice variant of Nrxn1β, and largely reverses the *cis*-Nrxn1β-mediated inhibition of trans-Nrxn1b/Nlgn2 interactions. ***A.*** Representative images of HEK293 cells transfected with Flag-tagged Nrxn1β^SS4+^, Nrxn1β^SS4−^, or a mock control and sequentially incubated for 1.5 h first with HA-Cbln1 or Nlgn1-ECD-Fc, and second with vehicle, Nlgn1-ECD-Fc, or HA-Cbln1 as indicated. Binding of HA-Cbln1 and Nlgn1-ECD-Fc to the surface-exposed Nrxn1β was visualized by immunocytochemistry in non-permeabilized cells. ***B.*** Demonstration that Cbln1 prevents and competes off binding of Nlgn1-ECD-Fc to Nrxn1β^SS4+^ but not to Nrxn1β^SS4−^ (summary graphs of surface binding of HA-Cbln1 (green) or Nlgn1-ECD-Fc (blue) to Nrxn1β^SS4+^ or Nrxn1β^SS4−^). ***C.*** Representative images of freestyle HEK cells transfected with EGFP and mCherry together with respective synaptic organizer molecules. Nrxn1β* denotes Nrxn1β with Nlgn binding mutations (G155V, T156A). ***D.*** Summary of Pearson’s coefficients across different conditions. All numerical data are means ± SEM. Sample size is indicated by numbers of cells/culture batches in B in which fluorescence intensity from cells in each batch was averaged as one data point; and n=4 batches of culture were used in C-D. Statistical analyses were performed by two-way ANOVA with Bonferroni’s multiple comparison test in D (****: p<0.0001).

The binding data suggests that Cbln1 could prevent a *cis-*interaction of neurexins with neuroligins, and thereby function in reciprocal synapses by relieving the inhibition of mitral cell neuroligins that is induced by *cis*-binding of neurexins (Fig. 1B). To test this hypothesis, we examined the inhibitory effect of *cis*-neurexin/neuroligin interactions on *trans*-neurexin/neuroligin interactions, and tested the possible role of Cbln1 in blocking this inhibition (Fig. 7C). We reconstituted *cis*- vs. *trans*-neurexin/neuroligin interactions using cell-aggregation assays with freestyle HEK293 cells that co-express Nrxn1β or Nlgn2 to mimic the expression observed in mitral cells (Fig. 1E, 1H). We then tested the effect of the *cis*-expression of Nrxn1β^SS4+^ or Nrxn1β^SS4−^ with Nlgn2 on the *trans*-Nrxn1β/Ngln2 interaction in the absence and presence of Cbln1 (Fig. 7C). Although Nlgn2-expressing cells formed large cell aggregates with Nrxn1β-expressing cells, these aggregates were suppressed when Nrxn1β, independent of SS4, was also co-expressed in *cis* with Nlgn2. Hence, the *cis* Nrxn1β-Nlgn2 interaction impaired the *trans*-Nrxn1β/Nlgn2 interaction without completely abolishing it (Fig. 7D). As a control, we co-expressed in *cis* Nrxn1β mutants that are unable to bind to neuroligins (Nrxn1β^SS4−^* or Nrxn1β^SS4+^*). These mutants contain G155V and T156A substitutions that block neuroligin binding (Reissner et al., 2008). *cis*-Expression of Nrxn1β mutants, in contrast to wild-type Nrxn1β, had no effect on the *trans*-Nrxn1β/Nlgn2 interaction (Fig. 7D).

Importantly, when we co-expressed Nlgn2 not only with Nrxn1β^SS4+^ or Nrxn1β^SS4−^, but also with Cbln1, the co-expressed Cbln1 reversed the inhibition of the *trans*-Nrxn1β/Nlgn2 interaction that was produced by *cis*-expression of Nrxn1β^SS4+^ but not the inhibition that was produced by *cis*-expression of Nrxn1β^SS4−^ (Fig. 7D). These data demonstrate that a *cis*-neurexin inhibits *trans*-neurexin/neuroligin interactions, and that this inhibition is alleviated when a cerebellin is co-expressed in *cis* with the neurexin and neuroligin, as long as the isoform of neurexin used is capable of actually binding to the cerebellin. Hence, the two antiparallel neurexin signaling processes can interfere with each other, and Cbln1 can mediate their functional compartmentalization, underlying the design principles for the reciprocal synapses.

### Cbln1 in mitral cells functions to compartmentalize neurexin signaling in reciprocal dendro-dendritic synapses

Our data thus suggest a molecular logic of reciprocal synapses whereby Cbln1 blocks *cis*-interactions of neurexins with neuroligins and thereby enables antiparallel *trans*-interactions of neurexins and neuroligins. Via this mechanism, Cbln1 functionally separates pre- and postsynaptic signaling compartments. An inherent prediction of this hypothesis is that postsynaptic neurexins in mitral cells are not essential for GC→MC synaptic transmission, but rather function as presynaptic neurexins for the mitral cell→granule cell synapses. This hypothesis also implies that the conversion of postsynaptic SS4+ neurexins into SS4− neurexins impairs GC→MC synaptic transmission only because it prevents Cbln1 from blocking the *cis*-interactions of postsynaptic neurexins with neuroligins. To test this hypothesis directly, we deleted all neurexins (except for Nrxn1γ that does not bind to either Cbln1 or neuroligins [Sterky et al., 2017]) from mitral cells by injection of rAAV2-retro’s that co-express EGFP with Cre or ΔCre (as a control) into the piriform cortex of *Nrxn123* triple conditional KO mice (Chen et al., 2017), and analyzed GC→MC synapses in acute slices by mitral cell recordings (Fig. 8A). As predicted by the hypothesis, the postsynaptic deletion of neurexins had no effect on synaptic transmission, and thus postsynaptic neurexins are functionally compartmentalized from the adjacent antiparallel neurexin-neuroligin signaling process (Fig. 8B-D; Fig. S7A-E).

**Figure 8.**
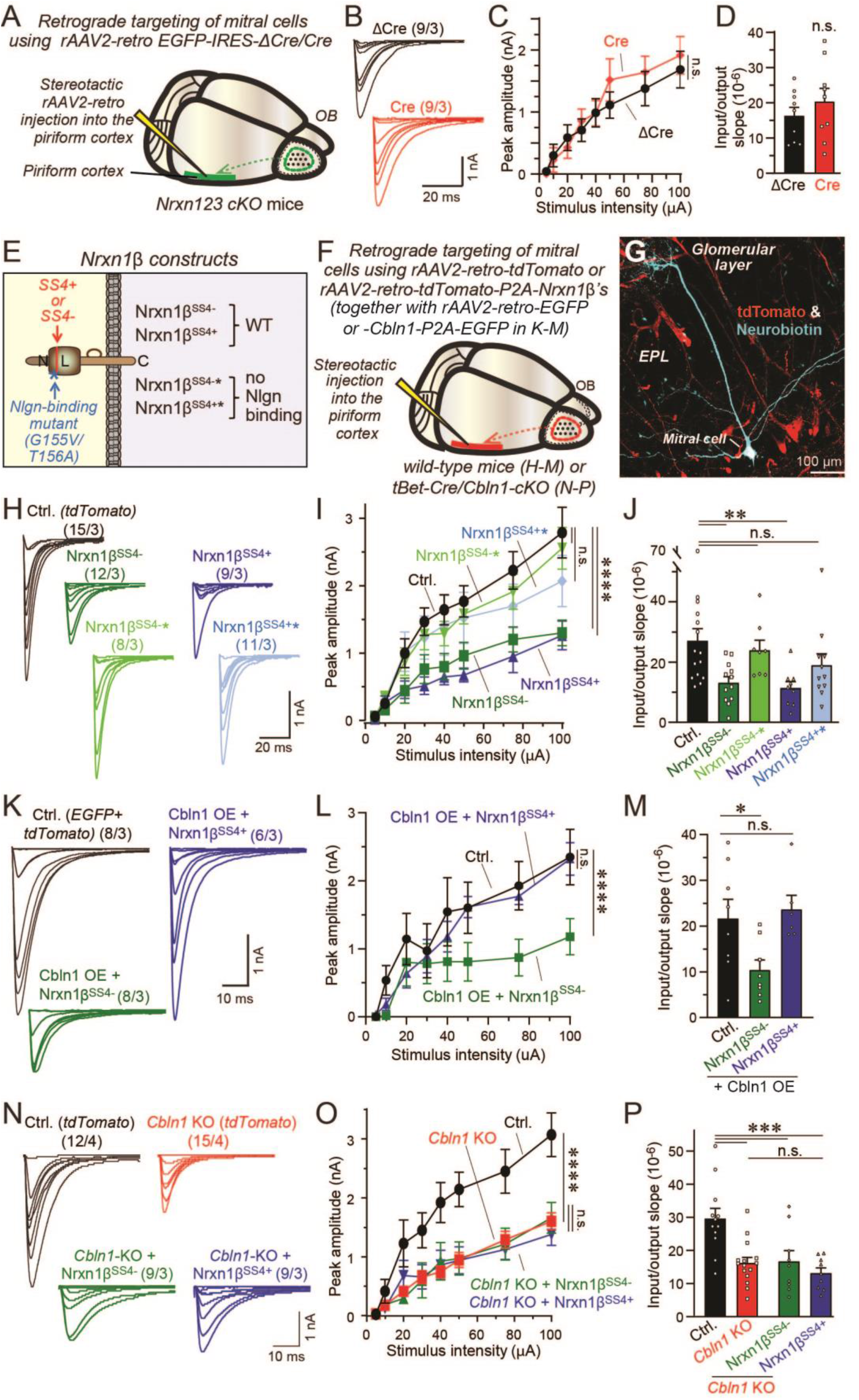
Postsynaptic neurexins in mitral cells are not essential for GC→MC synaptic transmission but suppress GC→MC synaptic transmission by *cis*- inhibition of postsynaptic neuroligins that is alleviated by Cbln1 binding. ***A***. Experimental strategy for the postsynaptic deletion of all neurexins in mitral cells. rAAV2-retro co-expressing EGFP with Cre or ΔCre (as a control) were injected into the piriform cortex of *Nrxn123* triple cKO mice to target mitral cells. ***B-D***. Postsynaptic deletion of neurexins does not impair GC→MC synaptic transmission (B, representative traces of evoked GC→MC IPSCs; D & E, input/output curve of the GC→MC IPSC peak amplitude (D) and summary graph of the input/output slope determined in individual experiments). ***E.*** Schematic of the Nrxn1β domain structure and overview of the Nrxn1b constructs used for overexpression experiments in mitral cells in panels F-M. ***F.*** Experimental strategy for the postsynaptic overexpression of Nrxn1β constructs in mitral cells. rAAV2-retro co-expressing tdTomato with the indicated Nrxn1β proteins (or without a Nrxn1β in control groups; or with EGFP/Cbln1-P2A-EGFP in K-M) were injected into the piriform cortex of wild-type mice or of mitral cell-specific *Cbln1* KO mice in which Cbln1 was deleted from mitral cells of *Cbln1* cKO mice using the *tBet-Cre* driver line. ***G.*** Representative image of a recorded mitral cell filled with neurobiotin and tdTomato (expressed in mitral cells via the rAAV2-retro infection). ***H-J***. Postsynaptic Nrxn1β overexpression in mitral cells inhibits GC→MC synaptic transmission for the wild-type Nrxn1β ^SS4+^ and Nrxn1β ^SS4−^ but not for mutant Nrxn1β^SS4+^* and Nrxn1β^SS4−^* that are unable to bind to neuroligins. (H, representative traces of evoked GC→MC IPSCs; I & J, input/output curves of the peak amplitudes of GC→MC IPSCs (I) and summary graph of the input/output curve slopes (J)). Note that Nrxn1β^SS4+^ likely inhibits despite the presence of Cbln1 because Cbln1 levels are insufficient to saturate binding of Nrxn1β^SS4+^. ***K-M***. The inhibition of GC→MC synaptic transmission by overexpressed Nrxn1β^SS4+^ but not Nrxn1β^SS4−^ is occluded by Cbln1 overexpression (OE) (K, representative traces of evoked GC→MC IPSCs; L & M, input/output curves of the peak amplitudes of GC→MC IPSCs (L) and summary graph of the input/output curve slopes (M)). ***N-P***. The inhibition of GC→MC synaptic transmission by overexpressed Nrxn1β^SS4+^ and Nrxn1β^SS4−^ is occluded by prior deletion of *Cbln1*, placing Nrxn1β and Cbln1 into the same pathway (N, representative traces of evoked GC→MC IPSCs; O & P, input/output curves of the peak amplitudes of GC→MC IPSCs (O) and summary graph of the input/output curve slopes (P)). All numerical data are means ± SEM (numbers of cells/mice analyzed are indicated above the sample traces). Statistical analyses were performed by *Student’s* t-test in D, by one-way ANOVA with Bonferroni’s multiple comparison test in J, M and P, and by two-way ANOVA with Bonferroni’s multiple comparison test in C, I, L and O (*, p<0.05; **, p<0.01; ***, p<0.001; ****, p<0.0001).

In a final test of this hypothesis, we overexpressed in mitral cells either wild-type Nrxn1β^SS4−^ and Nrxn1β^SS4+^ or mutant Nrxn1β^SS4−^* and Nrxn1β^SS4+^* that are unable to bind to neuroligins (Reissner et al., 2008; Fig. 7C-D, 8E). We again injected rAAV2-retro’s into the piriform cortex to selectively express the various neurexins in mitral cells of the OB (Fig. 8F-G). Overexpression of Nrxn1β^SS4−^ and Nrxn1β^SS4+^ inhibited GC→MC synaptic transmission as measured by evoked IPSCs, whereas overexpression of mutant Nrxn1β^SS4−^* and Nrxn1β^SS4+^* had no effect (Fig. 8H-J; Fig. S7F-L). In these experiments, even Nrxn1β^SS4+^ was inhibitory despite its binding to Cbln1 probably because the Nrxn1β^SS4+^ overexpression levels exceeded those of endogenous Cbln1. Hence, we tested the effect of co-overexpression of Cbln1 and Nrxn1β^SS4−^ (or Nrxn1β^SS4+^). Not surprisingly, Cbln1 overexpression in mitral cells likely saturates binding of Nrxn1β^SS4+^ and hence occludes its dominant negative effect, but not for Nrxn1β^SS4−^ (Fig. 8K-M; Fig. S7M-S). To test the physiological relevance of Cbln1 in the *cis* inhibition by Nrxn1β, we overexpressed Nrxn1β^SS4−^ and Nrxn1β^SS4+^ on the background of the *Cbln1* deletion and observed no additive effects (Fig. 8N-P; Fig. S7T-Z). Thus, Cbln1 functionally compartmentalizes two antiparallel neurexin signaling processes by preventing *cis*- neurexin/neuroligin interactions at the reciprocal dendro-dendritic synapses in the OB.

## DISCUSSION

Reciprocal dendro-dendritic synapses are present in many brain regions, and are particularly important in the OB for olfactory information processing (Shepherd et al., 2020). Reciprocal dendro-dendritic synapses pose a unique cell-biological problem in that they comprise two antiparallel synapses on adjacent dendrites, each of which forms pre- and post-synaptic specializations in the same subcellular compartment (Fig. 1A-B; Rall et al., 1966). Reciprocal synapses constitute a microcircuit that computes information without the involvement of action potentials, is highly plastic, and contributes to olfactory learning (Jahr & Nicoll, 1980; Isaacson & Strowbridge, 1998; Liu et al., 2017). Previous studies elucidated the detailed electrophysiological features of reciprocal synapses in the OB (Chen, Xiong & Shepherd, 2000; Halabisky et al., 2000; Isaacson, 2001; Balu et al., 2007; Gao & Strowbridge, 2009; Pressler & Strowbridge, 2017). However, the molecular mechanisms that organize dendro-dendritic reciprocal synapses remain unclear. In particular, the cell-biological conundrum of how functional pre- and post-synaptic specializations co-exist within the same subcellular compartment is unsolved (Fig. 1B). The lack of physical compartmentalization in reciprocal synapses implies that pre- and postsynaptic specializations share signaling molecules, creating the potential for ‘mixed signals’. Both mitral cells and granule cells express high levels of neurexins (Ullrich, Ushkaryov & Südhof, 1995; Uchigashima et al., 2019) and of the neurexin ligands cerebellins and neuroligins (Fig. 1F-G). The question of how multiple molecular machineries involving neurexins and their different ligands operate in parallel in reciprocal synapses is crucial for understanding their functional architecture (Fig. 1B).

Our data reveal a basic design principle in the organization of reciprocal dendro-dendritic synapses that explains how *cis*- and *trans*-interactions of synaptic adhesion molecules can be segregated. We show that in mitral cells, Cbln1 is essential for GC→MC synaptic transmission because Cbln1 blocks the *cis*-interaction of postsynaptic mitral-cell neurexins with neuroligins. When *Cbln1* is deleted and this *cis*-interaction is enabled, the *trans*-synaptic binding of postsynaptic mitral-cell neuroligins with presynaptic granule-cell neurexins is abolished, and GC→MC synaptic transmission is impaired.

The evidence for this conclusion is as follows. First, we show that in mitral cells, Cbln1 is essential for the organization of GC→MC synapses. Cbln1 is not required for the establishment or maintenance of GC→MC synapses or for regulating the presynaptic release probability, but instead enables a physiological postsynaptic GABA_A_-receptor response (Fig. 2-4). Second, we demonstrate that neuroligins are also essential for a normal postsynaptic GABA_A_-receptor response without a role in the presynaptic release probability (Fig. 5). Third, we showed that mitral cells express almost only SS4+ variants of neurexins that bind to Cbln1, and that converting mitral-cell SS4+ neurexins into SS4− neurexins impairs GC→MC synapses in the same manner as the Cbln1 and the neuroligin deletions (Fig 6). Fourth, we describe that Cbln1 displaces Nlgn1 from Nrxn1β on the cell surface as long as Nrxn1β is expressed as an SS4+ variant that binds Cbln1, that Nrxn1β–when expressed in *cis* with Nlgn2– suppresses the *trans*-interaction of Nlgn2 with Nrxn1β present on another cell, and that this suppression is alleviated by co-expression of Cbln1 (Fig.7). Finally, we documented that postsynaptic neurexins in mitral cells are not essential for GC→MC synaptic transmission, but that overexpression of postsynaptic Nrxn1β impairs GC→MC synapses as long as the overexpressed Nrxn1β is able to bind to neuroligins (Fig. 8).

Together, these data reveal that postsynaptic mitral-cell Cbln1 organizes GC→MC synapses by preventing inhibitory *cis*-interactions of mitral-cell neurexins and neuroligins, thereby functionally compartmentalizing the antiparallel *trans*-neurexin interactions. As an alternative explanation, it seems unlikely that postsynaptic Cbln1 acts on presynaptic granule cell neurexins because conversion of SS4+ neurexins into SS4− neurexins (that don’t bind Cbln1; Uemura et al., 2011) did not impair GC→MC synaptic transmission (Fig. S5J-S). The fact that the postsynaptic pan-neurexin deletion also did not further increase GC→MC synaptic transmission rules out a postsynaptic function of neurexins (Fig. 8A-E). Thus, to the best of our knowledge no plausible alternative hypotheses account for our results.

Our data raise several new questions about reciprocal synapses. Does Cbln1 have an additional role in mitral cells, other than functionally compartmentalizing anti-parallel neurexin signaling processes in dendro-dendritic synapses? The fact that the postsynaptic *Clbn1* deletion not only reduced the mIPSC frequency, but also the mEPSC frequency, suggests that Cbln1 functions also at excitatory input synapses in mitral cells (Fig 2I-K). Moreover, it remains an open question whether Cbln1 performs additional *trans*-synaptic signaling at MC→GC synapses through GluD1 and GluD2 in granule cells. Furthermore, we showed earlier that the presynaptic deletion of *Nrxn3* in granule cells decreases the release probability at GC→MC synapses (Aoto et al., 2015). Since the postsynaptic neuroligin or Cbln1 deletions do not cause such a phenotype, other neurexin ligands must be involved. Clearly our data do not rule out the possibility of multiple parallel neurexin signaling pathways in a single synapse. After all we only examined cerebellins and neuroligins, but LRRTMs, dystroglycan, neurexophilins, and other potential neurexin ligands such as calsyntenins are likely present in mitral cells (reviewed in Südhof, 2017). In addition, non-neurexin pathways involving neurexin ligands – such as those mediated by MDGAs that bind to neuroligins (Connor et al., 2016 and 2017; Elegheert et al., 2017), are likely important, as are other synaptic adhesion systems such as those effected by adhesion GPCRs (Südhof, 2021). Using reciprocal synapses as a model synapse, we are only beginning to understand how multiple molecular machineries operate in parallel to shape synapse dynamics.

Finally, looking beyond reciprocal synapses in the OB, our work highlights the importance of multiple *trans*-synaptic signaling pathways at a given synapse. Most molecular and genetic studies on synaptic organizer molecules focus on components of one given *trans*-synaptic complex at a time (Südhof, 2021; Yuzaki, 2018). However, multiple molecular machineries likely operate in parallel, with or without lateral interactions, at all synapses to mediate synapse formation and to regulate their properties. *Trans*-synaptic molecular networks are likely organized similar to logic gates, with divergent and convergent signaling pathways and AND/OR/NOT decision points. Dissecting these functional networks is a painstakingly slow process that requires non-scalable, technically demanding methods, such as high-resolution electrophysiology or imaging. The molecular design principle for the reciprocal synapses shown here provides a first step towards such a dissection and represents the first analysis of the relation of multiple synaptic adhesion interactions, but much more remains to be done.

## ACKNOWLEDGEMENTS

This work was supported by a Stanford Interdisciplinary Graduate Fellowship (to C.Y.W.) and a grant from the National Institutes of Mental Health (MH052804 to T.C.S.).

## AUTHOR CONTRIBUTIONS

Conceptualization, C.Y.W. and T.C.S.; Methodology, C.Y.W., J.H.T., K.L-A, S-J.L., X.L. and T.C.S.; Formal analysis, C.Y.W., J.H.T., K.L-A, S-J.L., X.L. and T.C.S.; Investigation, C.Y.W., J.H.T., K.L-A, S-J.L. and X.L.; Writing – Original Draft, C.Y.W. and T.C.S.; Writing – Review & Editing, C.Y.W., J.H.T., K.L-A, S-J.L., X.L. and T.C.S.; Project Administration, C.Y.W. and T.C.S.

## MATERIALS and METHODS

### KEY RESOURCE TABLE

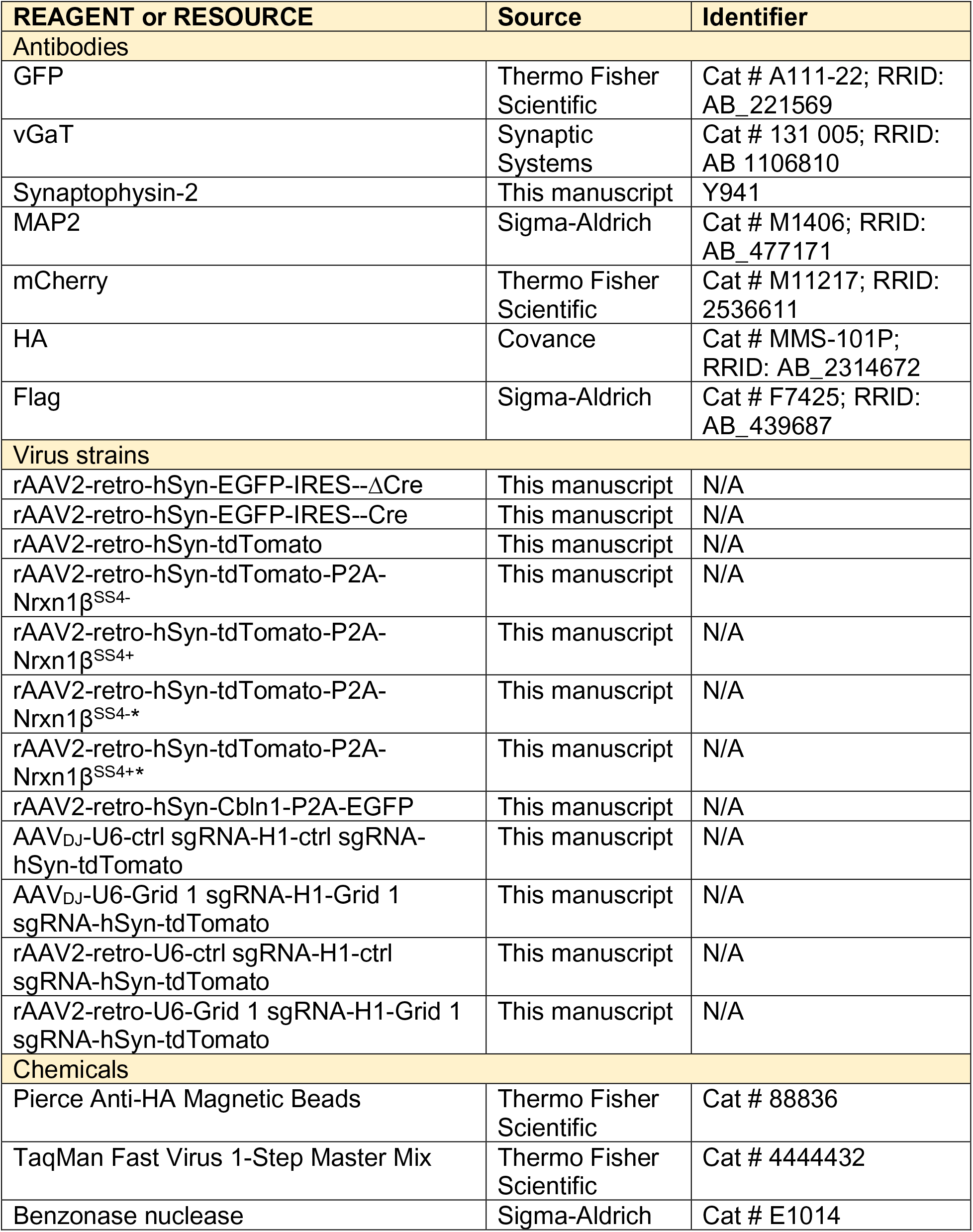
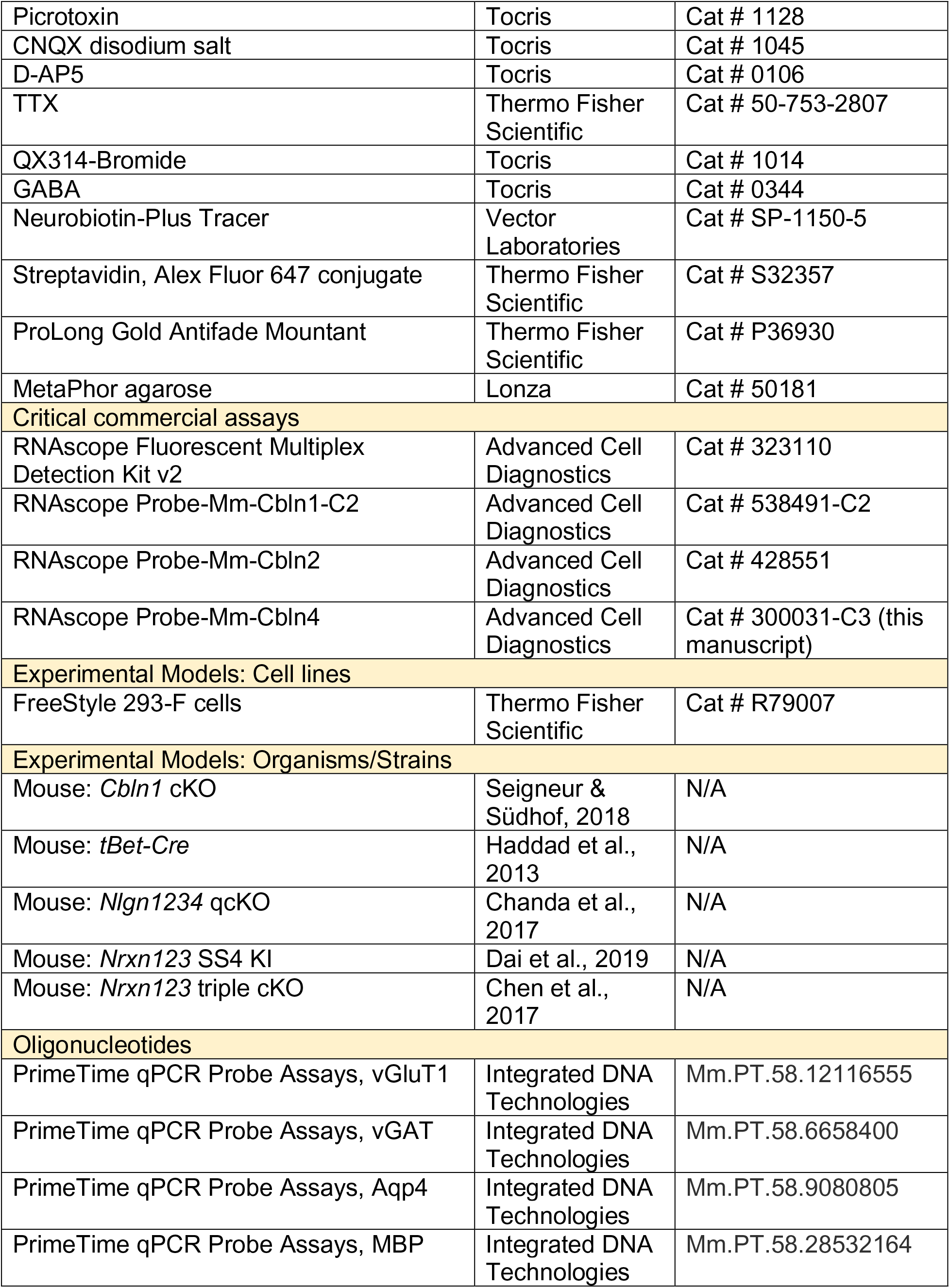
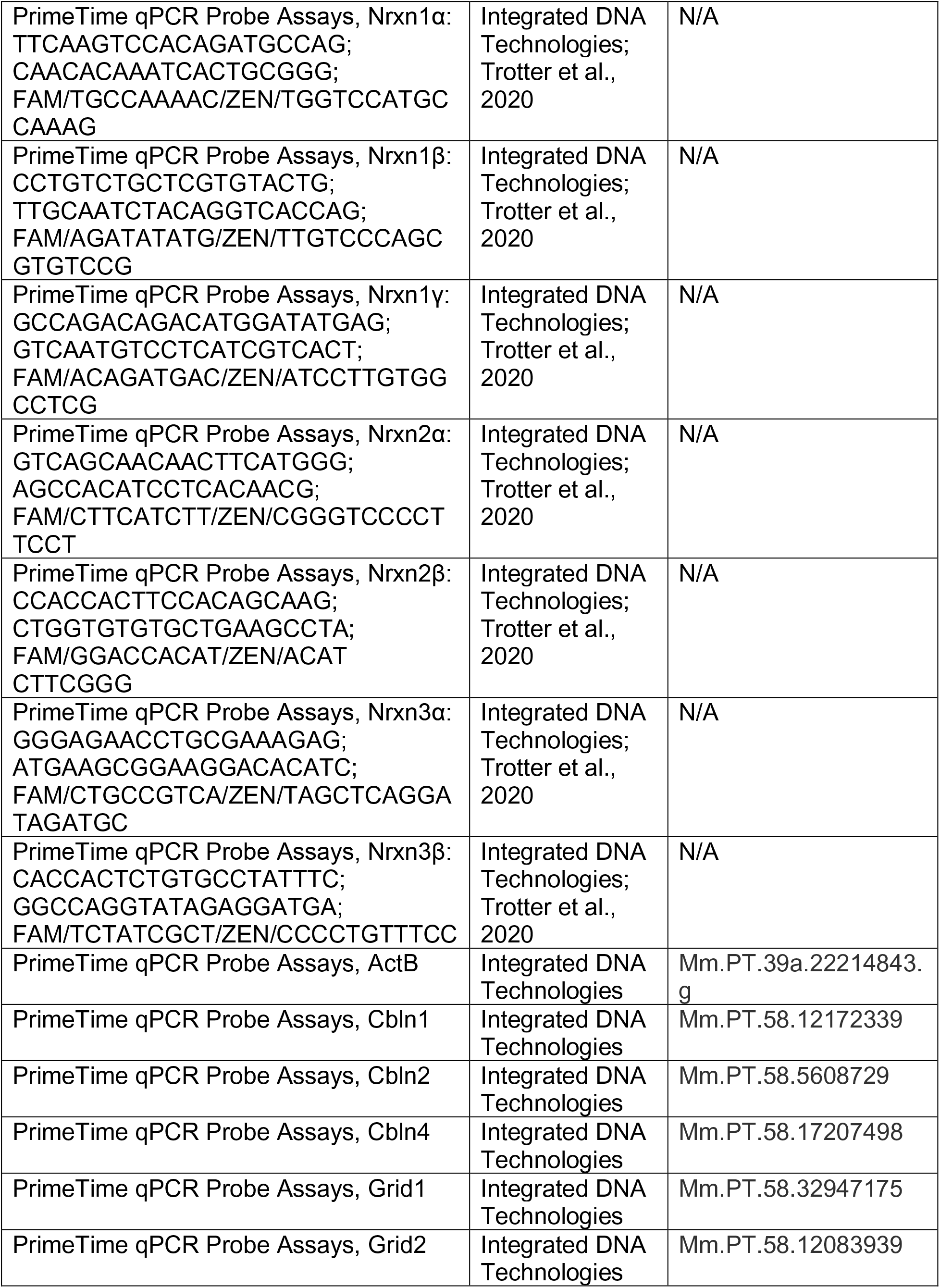
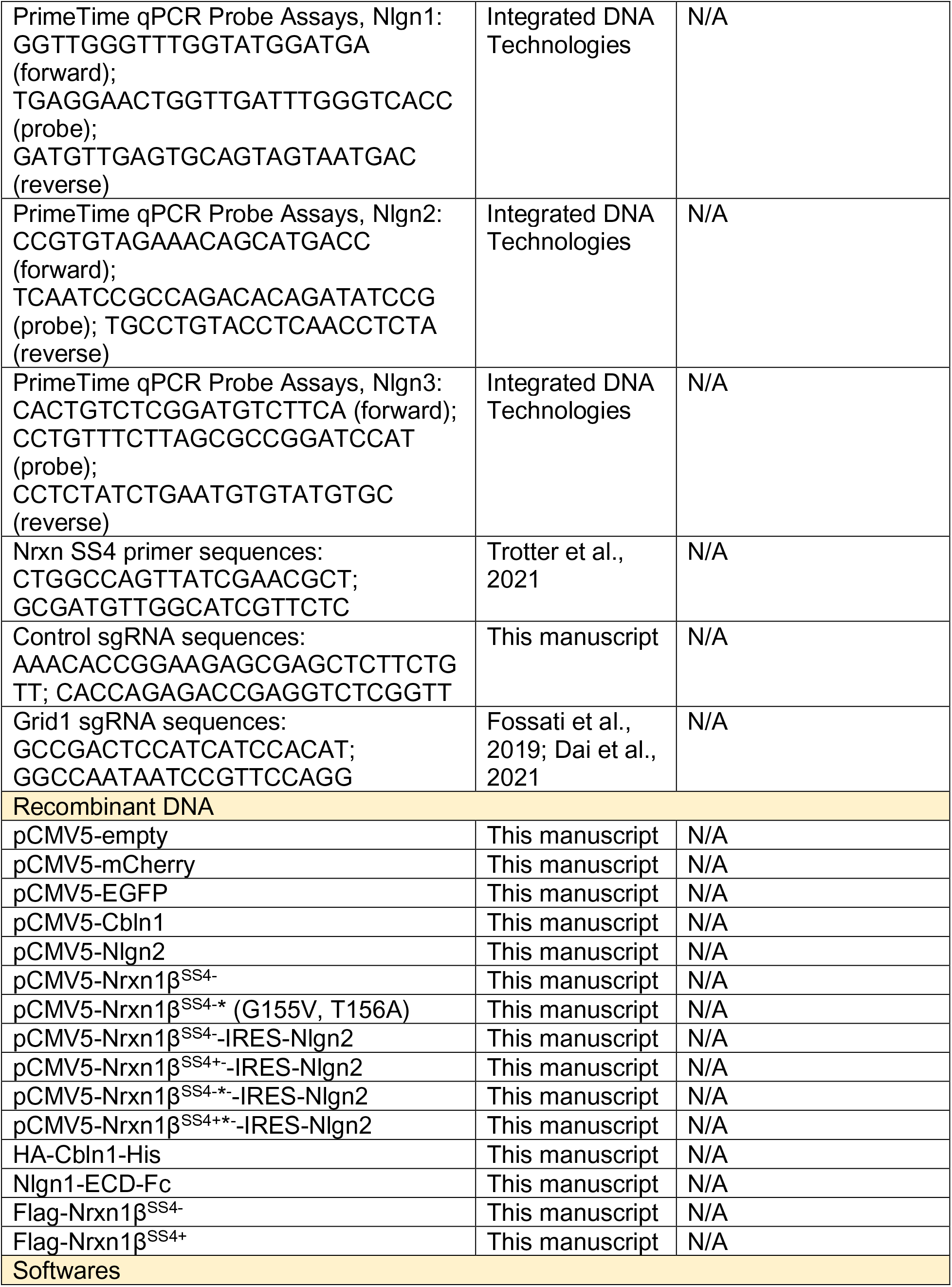
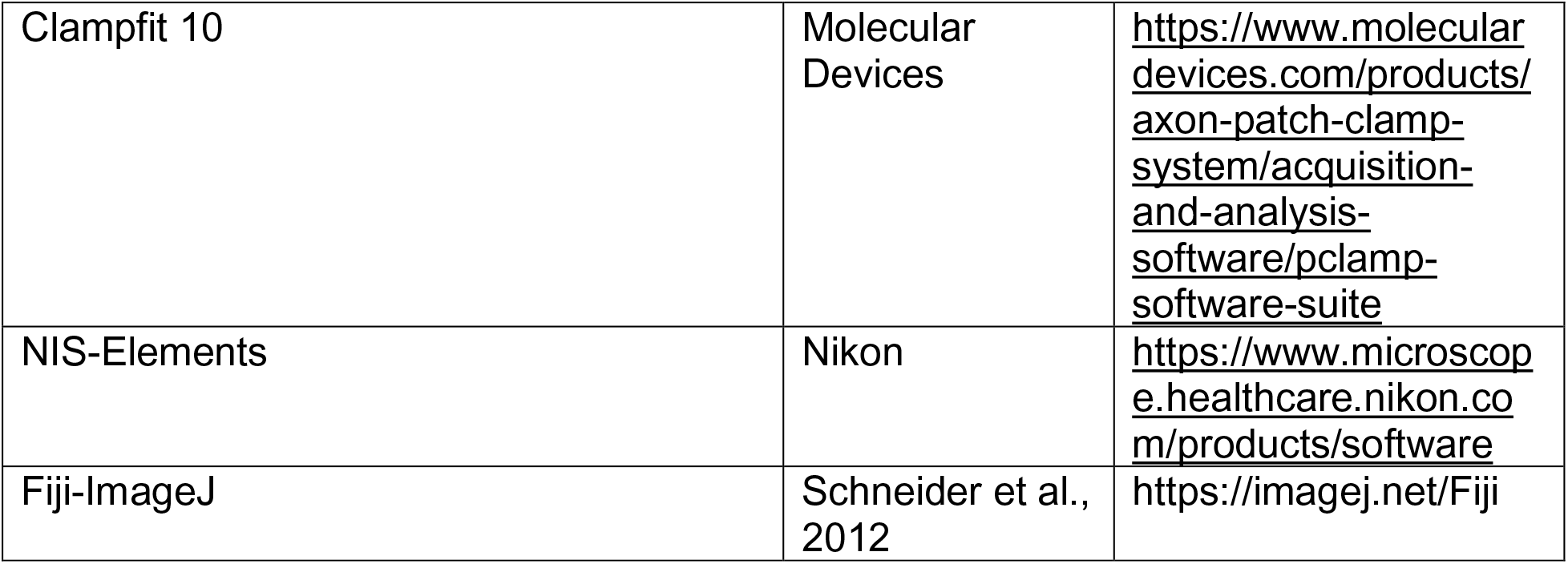

### RESOURCE AVAILABILITY

#### Lead Contact

Further information and requests for resources and reagents should be directed to and will be fulfilled by the Lead Contact, Thomas C. Südhof.

#### Materials Availability

Plasmids generated in this study for virus production and *in vitro* experiments are available upon reasonable requests.

#### Data and Code Availability

No code is generated in this study. Raw data are available upon reasonable request.

### EXPERIMENTAL MODEL AND SUBJECT DETAILS

For mitral/tufted-cell-specific *Cbln1* deletion, *Cbln1* homozygous cKO mice were crossed with *Cbln1* homozygous cKO mice with *tBet-Cre* allele (Seigneur & Südhof, 2018; Haddad et al., 2013). *Nlgn1234* qcKO mice (Chanda et al., 2017), *Nrxn123* SS4 cKI mice (Dai et al., 2019) and *Nrxn123* cKO mice (Chen et al., 2017) were generated as described in respective previous studies. Mice were group-housed (maximum of five mice per cage) and maintained on a 12 h light–dark cycle (7 am to 7 pm, light), with access to food and water *ad libitum*. Age- and gender-matched littermates were used for experiments, except for the Nrxn1β overexpression experiments in Fig.8 where the number of mice required made it impossible and only age-matched littermates were used. All experiments involving animals were approved by the Stanford Animal Use Committees (IACUC and APLAC).

### METHOD DETAILS

#### Purification of olfactory bulb mRNA

Mice were euthanized using isoflurane and decapacitated. The olfactory bulbs were quickly dissected and snap-frozen in liquid nitrogen or dry ice and transferred to −80°C storage until processing. The specimen was subjected to RNA extraction using the QIAGEN RNeasy Micro kit.

#### Purification of mitral/tufted-cell-specific mRNA

Ribotag mice were crossed with *tBet-Cre* mice (Sanz et al., 2009; Haddad et al., 2013). After olfactory bulb extraction described above, the frozen bulbs were partially thawed in fresh homogenization buffer at 10% weight/volume and Dounce homogenized. Homogenates underwent centrifugation and 10% of the supernatant was used as input. The remaining supernatant was incubated with pre-washed anti-HA magnetic beads (Thermo) overnight at 4°C. The beads were washed three time with a high-salt buffer followed by elution with RLT lysis buffer containing 2-mercaptoethanol. The sample and the input were then subjected to mRNA extraction described above.

#### Quantitative real-time PCR

Quantitative RT-PCR was performed in triplicates for each condition. 20 ng RNA was used each reaction, in conjunction with TaqMan Fast Virus 1-Step Master Mix (Thermo) and gene-specific qRT-PCR probes (IDT).

#### Electrophysiology

*Cbln1* cKO mice with or without *tBet-Cre* alleles were patched between P21 and P28. For viral manipulations, mice were injected at P16-P21. Two to three weeks after viral injection, mice were anesthetized via isoflurane inhalation and brains were quickly removed. The brain was sliced in ice-cold oxygenated (95% O_2_ and 5% CO_2_) cutting solution (228 mM sucrose, 11 mM glucose, 26 mM NaHCO_3_, 1 mM NaH_2_PO_4_, 2.5 mM KCl, 7 mM MgSO_4_, and 0.5 mM CaCl_2_). Horizontal sections (300 µm thickness) were obtained using a vibratome and placed in oxygenated artificial cerebrospinal fluid (ACSF; 119 mM NaCl, 2.5 mM KCl, 1 mM NaH_2_PO_4_, 1.3 mM MgSO_4_, 26 mM NaHCO_3_, 10 mM glucose, and 2.5 mM CaCl_2_) at 32°C for 30 min. Slices were allowed to recover at room temperature for an additional 30 min. The recording chamber was temperature controlled and set to 32°C, and ACSF was perfused at 1 mL/min. The internal solution for whole-cell patch clamp contained 135 mM CsCl, 10 mM HEPES, 1 mM EGTA, 1 mM Na-GTP and 4 mM Mg-ATP pHed to 7.4. 10mM QX314-bromide was added for evoked recordings. 0.2% neurobiotin (VectorLab) was included for morphological reconstruction. The pipette resistance ranged from 1.8 to 2.5 MΩ. Mitral cells were identified in the mitral cell layer, and access resistance was under 10 MΩ throughout the experiment. 1µM TTX (Tocris) and 100µM picrotoxin (Tocris) were included in the bath solution for mEPSC. 1µM TTX (Tocris), 20µM CNQX (Tocris) and 50µM D-AP5 (Tocris) were included in the bath for mIPSC and GABA puffing. 20µM CNQX (Tocris) and 50µM D-AP5 (Tocris) were included in the bath for evoked IPSC recordings. GC→MC evoked IPSCs (eIPSCs) were recorded at −70 mV with a concentric bipolar electrode placed directly below the mitral cell with constant distance roughly at the junction between internal plexiform layer and granule cell layer, 30 µm below the surface of the slice. PGC→MC eIPSCs were recorded by placing the concentric bipolar electrode on the glomerulus containing the dendrites of the recorded mitral cell, which were visualized by including Alexa-488 in the internal solution. The same concentric bipolar electrodes were used for each experiment. However, different concentric bipolar electrodes were used across different experiments as they wear out; and this variability might cause the differences in absolute value of eIPSC observed. For GABA puffing, 10 µM GABA (Tocris) dissolved in the bath solution were loaded in glass pipette with resistance of 1.8-2.5 MΩ and connected to the Picropritzer. The GABA-containing pipette was placed near the mitral-cell soma with constant distance. GABA puff was delivered with 10 psi of various durations. All recordings were analyzed in Clampfit after applying a 500 Hz Gaussian filter. Rise time was defined as the time from 10% to 90% of peak amplitude, and decay time was defined as the time from 90% to 10% of peak amplitude. The experimenter was blind to the treatment groups during recordings and analysis.

#### AAV preparation

The adeno-associated virus (AAV) serotype used in this study was AAV-DJ, as well as rAAV2-retro for retrograde tracing experiments (Tervo et al., 2016). HEK293T cells were transfected with the AAV vector, the helper plasmid and the serotype-specific capsid plasmids by using calcium phosphate. Cells were harvested 72 hours post-transfection. Nuclei were lysed, were treated with Benzonase nuclease (Sigma-Aldrich, cat # E1014) and underwent iodixanol gradient ultracentrifugation (3 h at 65,000 rpm using a S80AT3 rotor). AAV were then concentrated and dialyzed in minimal essential media (MEM).

#### Stereotactic injections

Mice were prepared for stereotactic injections using standard procedures approved by the Stanford University Administrative Panel on Laboratory Animal Care. For anesthesia, the stock solution was made of dissolving 5 g tribromoethanol into 5 mL T-amyl alcohol, and further diluted 80 folds into PBS to make the working solution. 0.2 mL working solution (Avertin) per 10 grams body weight of mouse was used for anesthesia before mounting the mouse in the stereotax. The coordinates (AP/ML/DV from Bregma) and volumes for the intercranial injections are as follows: (1) OB: +4.3/±0.85/−1.7 and +5.3/±0.6/−1.5 with 1.0 uL virus and (2) Piriform cortex: −0.7/±3.7/−4.75 with 0.75 uL virus. The reference point on DV-axis for OB injection was on the surface of OB, while the reference point on DV-axis for piriform cortex injection was on the skull surface at AP/ML coordinate of −0.7/0.0.

#### Behavioral experiments

All behaviors were assessed in adult mice 2-3 months ago after viral injections. Only male mice were used for the three-chamber sociability test.

##### Open field test

Mice were individually placed in a 40 x 40 x 40 cm^3^ white plastic chamber in a well-lit room and allowed to move freely for 10 min. Locomotor and exploratory behaviors were recorded using a Viewer III tracking system (Biobserve). Total distance traveled and time spent in the 20 x 20 cm^2^ center of the square were quantified.

##### Three-chamber sociability test

A transparent three-chamber apparatus (60 x 30 x 30 cm^3^ per chamber) was used for sociability tests. For habituation, the subject was placed in the center chamber and allowed to explore the entire apparatus for 5 min. The peripheral chamber (left or right) in which the mouse spent more time was designated the preferred side. Subsequently, a stranger juvenile *Cbln1* cKO mouse was placed underneath an upside-down black wire-mesh cup within the non-preferred chamber. The subject mouse was allowed to explore all three chambers freely for 10 min, while being recorded using the Viewer III tracking system. Time spent in each chamber was quantified, and the sociability index was calculated by dividing the time spent in the stranger mouse-containing chamber by the time spent in the empty chamber.

##### Buried food-finding test

After 24 h of food deprivation, the subject was placed in a new cage containing 3 cm of bedding and a 5 g food pellet buried in a random corner of the cage. The time it took for the mouse to dig up the food pellet was recorded.

#### Single-molecule in situ hybridization

Wild-type CD-1 mice were euthanized with isoflurane at P30 followed by transcardial perfusion with ice-cold PBS. Brains were quickly dissected and embedded in Optimal Cutting Temperature (OCT) solution on dry ice. Sections with 12-µm thickness were sliced using a Leica cryostat 9CM3050-S) and mounted directly onto Superfrost Plus histological slides. Single-molecule fluorescence *in situ* hybridization for Cbln1/2/4 mRNA (probe cat # 538491-C2, # 428551 and # 300031-C3 respectively) was performed using the multiplex RNAscope platform (ACD) according to manufacturer’s instruction. Slides were mounted using ProLong Gold antifade mounting medium (Thermo, P36930) and imaged using Olympus VS120 slide scanner.

#### Immunohistochemistry

For synaptic marker staining, mice were anesthetized by avertin injection as described in stereotactic injection. Brains were extracted and post-fixed in 4% PFA for 1 h. After post-fixation, brains were washed three times with PBS and cryoprotected in 30% sucrose in PBS for 24-36 h. Brains were sectioned coronally at 40 µm using a cryostat. For morphological tracing after electrophysiology, the slices were incubated in 4% PFA for 15 minutes. For immunostaining, the slices were blocked and permeabilized in 5% normal goat serum and 0.3% Triton X-100-containing PBS for 1 h at room temperature. Primary antibodies against vGaT (SySy, 1:1000), synaptophysin-2 (homemade with code Y941, 1:1000), MAP2 (Sigma, 1:1000), GFP (Thermo, 1:1000), or mCherry (Thermo, 1:1000) were diluted in blocking buffer and brain slices were incubated with primary antibody overnight at 4°C. For neurobiotin tracing, Streptavedin conjugated with Alexa Fluor 647 (Thermo, 1:1000) was included in the primary antibody mix. After three 10 min washes in PBS, slices were incubated with secondary antibodies: Alexa Fluor-488, −555 or −647. Slices underwent three 10 min washes in PBS and were mounted on positively-charged glass slides, allowed to dry, and coverslipped.

#### Confocal microscopy

A Nikon A1RSi confocal microscope was used to acquire all images. Images are analyzed using Nikon analysis software. Within each set of experiments, the laser power, gain, offset, and pinhole size for each laser were kept constant. For quantification of synaptic puncta, z-stack images were obtained by at 0.5 µm intervals and three slices (1 µm thickness in total) with highest signal were maximally projected. Automated background subtraction was performed using a rolling ball algorithm with a 1 µm radius. The same threshold was applied to each set of experiments and puncta parameters were automatically obtained using the analysis software. For quantification in the olfactory bulb, the entire olfactory bulb was sliced. Five sections with 400 µm anterior-posterior distance spacing were quantified, and each data point is an average of all slices quantified. The images were taken at either medial or lateral olfactory bulb and were chosen randomly. All IHC data were collected and analyzed blindly.

#### Transmission electron microscopy

Three pairs of P28 *tBet-Cre/Cbln1* cKO mice and littermate controls without *tBet-Cre* allele were perfused with PBS followed by 4% PFS at 37°C. The brains were dissected out and post-fixed in 2.5% glutaraldehyde and 2% paraformaldehyde in 0.1M sodium cacodylate buffer (pH 7.4) overnight at 4°C. 200µm coronal vibratome sections of the olfactory bulb were collected in 0.1M cold cacodylate buffer on the next day. The slices were subject to post-fixation in 1% glutaraldehyde in 0.1M cacodylate buffer before shipping to Yale CCMI EM facility. Slices were then post-fixed in 1% OsO_4_, 0.8% potassium ferricyanide in 0.1M cacodylate buffer at RT for 1 hour. Specimens were then *en bloc* stained with 2% aqueous uranyl acetate for 45 min, dehydrated in a graded series of ethanol to 100%, substituted with propylene oxide and embedded in Embed 812 resin. Sample blocks were polymerized in an oven at 60°C overnight. Thin sections (60 nm) were cut by a Leica ultramicrotome (UC7) and post-stained with 2% uranyl acetate and lead citrate. Sections were examined with a FEI Tecnai transmission electro microscope at 80 kV of accelerating voltage, digital images were recorded with an Olympus Morada CCD camera and iTEM imaging software. Image analysis was performed using Fiji-ImageJ. Excitatory synapses from mitral cells were identified by the presence of asymmetric membrane thickening and at least two presynaptic vesicles. Inhibitory synapses onto mitral cells were identified by the presence of symmetric membrane thickening and at least two presynaptic elliptical vesicles as compared to excitatory synapses. Synapse density were quantified by randomly sampling mitral cell dendrites, quantifying the number of synapses along the dendrites and normalizing the number by the total length of mitral cell dendrites analyzed in each mouse. Bouton size, the number of vesicles per bouton, the average vesicle size, vesicle ellipticity and the average distance of vesicles to the active zone were quantified for inhibitory synapses onto mitral cells. All the parameters were averaged for each animal for statistical analysis. All EM data were acquired and analyzed blindly.

#### Junction-flanking PCR

A pair of primers (5’-CTGGCCAGTTATCGAACGCT-3’; 5’-GCGATGTTGGCATCGTTCTC-3’) annealing to constitutive exon sequences that flank splice site 4 exon were used to amplify *Nrxn123* mRNA transcripts with or without SS4. cDNA was firstly synthesized from equal amounts of immunoprecipitated mRNA from mitral/tufted cells and total input mRNA from the OB. Junction-flanking PCR was then performed with equal amount of cDNA from the two groups. The PCR products were separated on homemade MetaPhor agarose gel (Lonza) and stained with GelRed. Stained gel was imaged at sub-saturation using the ChemiDoc Gel Imaging System (Bio-Rad). Quantification was performed using Image Lab (Bio-Rad) or ImageStudioLite (LI-COR). Intensity values were normalized to the size of DNA products to control for intensity differences caused by different dye incorporation owing to varied DNA length.

#### Surface binding assay and immunocytochemistry

Soluble recombinant HA-Cbln1- His and Nlgn1-ECD-Fc were firstly prepared as follows. FreeStyle HEK 293F cells (Thermo) were transfected with plasmids expressing either HA-Cbln1-His or Nlgn1-ECD- Fc. The cell medium was collected 5 d after transfection. His-tagged Cbln1 were purified by Talon metal affinity resin (Clontech) and dialyzed against HBSS. Fc-tagged Nlgn1- ECD were purified by rProteinA Sepharose Fast Flow (GE Healthcare), eluted with 10mM glycine, pH 2.5 and dialyzed with HBSS. After obtaining soluble proteins, HEK293 cells were transfected with expression vectors for Flag-Nrxn1β with or without SS4. Vehicle, HA-Cbln1 (50 µg/mL) and Nlgn1-ECD-Fc (50 µg/mL) in HBSS with 2 mM CaCl_2_ and 1 mM MgCl_2_ was added to the transfected HEK293 cell sequentially with each treatment lasting 1.5 h at room temperature. After washing, the cells were fixed with 4% PFA and immunostained with mouse anti-HA (Covance, 1:1000) or rabbit anti-Flag (Sigma, 1:1000), followed by incubation with AlexFluor-647-conjugated anti-human Fc γ and species-specific AlexaFluor-488- or -546-conjugated secondary antibodies. Coverslips were mounted after three times of washing in PBS.

#### Cell aggregation assay

One batch of FreeStyle HEK 293F cells were transfected with pCMV5-EGFP together with pCMV5-empty, pCMV5-Nrxn1β^SS4−^ or pCMV5-Nrxn1β^SS4−^* (with Nlgn binding mutations G155V, T156A). Another batch of FreeStyle HEK 293F cells were transfected with pCMV5-mCherry together with pCMV5-empty and pCMV5 expression vectors containing Nlgn2. *Cis* co-expression of Nrxn1β and Nlgn2 was achieved by constructing plasmids co-expressing both using IRES. Cbln1 was co-expressed by the addition of a third plasmid pCMV5-Cbln1 during transfection. 48 hours after transfection, 1mL of HEK 293F cells from each batch was mixed together in a 12-well Corning Costar Not Treated Plate (Millipore) and incubated at 37°C with shaking on Orbi-Shaker (Benchmark) with 125 rpm for 2h. Three images for each well were taken under confocal microscopy. Images were thresholded using Otsu’s method and Pearson’s coefficient was calculated. Each data point represented averaged Pearson’s coefficient for the three images taken for one round of experiments from transfection to imaging.

### QUANTIFICATION AND STATISTICAL ANALYSIS

All experiments of electrophysiology, behavioral tests, immunohistochemistry analysis, electron microscopy and cell culture experiments were performed and analyzed blindly to the experimental condition. *Student’s* t test was used whenever the comparison is between two groups. The Kolmogorov-Smirnov test was used to analyze the cumulative curves. One-way ANOVA with Bonferroni’s multiple hypothesis correction was used for comparison among more than two groups. Two-way ANOVA with Bonferroni’s multiple hypothesis correction was used for comparison of multiple groups with multiple factors. The statistical test used for each experiment was specified in the figure legend. The “*n*” used for these analyses represents the number of mice for gene expression analysis, behavioral tests, immunohistochemistry analysis and electron microscopy analysis, the number of cells for electrophysiology and perisomatic synapse quantification and the number of culture batches for surface binding assay and cell aggregation assay, all of which have been specified in Figure Legends.

## REFERENCES

Ango, F., Gallo, N.B., and Van Aelst, L. (2021). Molecular mechanisms of axo-axonic innervation. Curr. Opin. Neurobiol. 69, 105–112.

Aoto, J., Földy, C., Ilcus, S.M.C., Tabuchi, K., and Südhof, T.C. (2015). Distinct circuit-dependent functions of presynaptic neurexin-3 at GABAergic and glutamatergic synapses. Nat. Neurosci. 18, 997–1007.

Araç, D., Boucard, A.A., Özkan, E., Strop, P., Newell, E., Südhof, T.C., and Brunger, A.T. (2007). Structures of Neuroligin-1 and the Neuroligin-1/Neurexin-1β Complex Reveal Specific Protein-Protein and Protein-Ca2+ Interactions. Neuron 56, 992–1003.

Balu, R., Pressler, R.T., and Strowbridge, B.W. (2007). Multiple modes of synaptic excitation of olfactory bulb granule cells. J Neurosci 27, 5621–5632.

Bergmann, M., Schuster, T., Grabs, D., Marquèze-Pouey, B., Betz, H., Traurig, H., Mayerhofer, A., and Gratzl, M. (1993). Synaptophysin and synaptoporin expression in the developing rat olfactory system. Dev. Brain Res. 74, 235–244.

Bolliger, M.F., Pei, J., Maxeiner, S., Boucard, A.A., Grishin, N. V., and Südhof, T.C. (2008). Unusually rapid evolution of Neuroligin-4 in mice. Proc. Natl. Acad. Sci. U. S. A. 105, 6421–6426.

Chanda, S., Hale, W.D., Zhang, B., Wernig, M., and Südhof, T.C. (2017). Unique versus redundant functions of neuroligin genes in shaping excitatory and inhibitory synapse properties. J. Neurosci. 37, 6816–6836.

Chen, L.Y., Jiang, M., Zhang, B., Gokce, O., and Südhof, T.C. (2017). Conditional Deletion of All Neurexins Defines Diversity of Essential Synaptic Organizer Functions for Neurexins. Neuron 94, 611–625.

Chen, W.R., Xiong, W., and Shepherd, G.M. (2000). Analysis of relations between NMDA receptors and GABA release at olfactory bulb reciprocal synapses. Neuron 25, 625–633.

Chen, X., Liu, H., Shim, A.H.R., Focia, P.J., and He, X. (2008). Structural basis for synaptic adhesion mediated by neuroligin-neurexin interactions. Nat. Struct. Mol. Biol. 15, 50–56.

Connor, S.A., Ammendrup-Johnsen, I., Chan, A.W., Kishimoto, Y., Murayama, C., Kurihara, N., Tada, A., Ge, Y., Lu, H., Yan, R., et al. (2016). Altered Cortical Dynamics and Cognitive Function upon Haploinsufficiency of the Autism-Linked Excitatory Synaptic Suppressor MDGA2. Neuron 91, 1052–1068.

Connor, S.A., Ammendrup-Johnsen, I., Kishimoto, Y., Karimi Tari, P., Cvetkovska, V., Harada, T., Ojima, D., Yamamoto, T., Wang, Y.T., and Craig, A.M. (2017). Loss of Synapse Repressor MDGA1 Enhances Perisomatic Inhibition, Confers Resistance to Network Excitation, and Impairs Cognitive Function. Cell Rep. 21, 3637–3645.

Cover, K.K., and Mathur, B.N. (2020). Axo-axonic synapses: Diversity in neural circuit function. J. Comp. Neurol. 1, 2391–2401.

Dai, J., Aoto, J., and Südhof, T.C. (2019). Alternative Splicing of Presynaptic Neurexins Differentially Controls Postsynaptic NMDA and AMPA Receptor Responses. Neuron 102, 993–1008.

Dai, J., Patzke, C., Liakath-Ali, K., Seigneur, E., and Südhof, T.C. (2021) GluD1, A signal transduction machine disguised as an ionotropic receptor. Nature, in press.

Elegheert, J., Cvetkovska, V., Clayton, A.J., Heroven, C., Vennekens, K.M., Smukowski, S.N., Regan, M.C., Jia, W., Smith, A.C., Furukawa, H., et al. (2017). Structural Mechanism for Modulation of Synaptic Neuroligin-Neurexin Signaling by MDGA Proteins. Neuron 95, 896–913.

Eyre, M.D., Antal, M., and Nusser, Z. (2008). Distinct deep short-axon cell subtypes of the main olfactory bulb provide novel intrabulbar and extrabulbar GABAergic connections. J. Neurosci. 28, 8217–8229.

Famiglietti, E. V. (1970). Dendro-dendritic synapses in the lateral geniculate nucleus of the cat. Brain Res. 20, 181–191.

Fossati, M., Assendorp, N., Gemin, O., Colasse, S., Dingli, F., Arras, G., Loew, D., and Charrier, C. (2019). Trans-Synaptic Signaling through the Glutamate Receptor Delta-1 Mediates Inhibitory Synapse Formation in Cortical Pyramidal Neurons. Neuron 104, 1081–1094.

Gao, Y., and Strowbridge, B.W. (2009). Long-term plasticity of excitatory inputs to granule cells in the rat olfactory bulb. Nat Neurosci 12, 731–733.

Haddad, R., Lanjuin, A., Madisen, L., Zeng, H., Murthy, V.N., and Uchida, N. (2013). Olfactory cortical neurons read out a relative time code in the olfactory bulb. Nat. Neurosci. 16, 949–957.

Halabisky, B., Friedman, D., Radojicic, M., and Strowbridge, B.W. (2000). Calcium Influx through NMDA Receptors Directly Evokes GABA Release in Olfactory Bulb Granule Cells. J. Neurosci. 20, 5124–5134.

Harding, B.N. (1971). Dendro-dendritic synapses, including reciprocal synapses, in the ventrolateral nucleus of the monkey thalamus. Brain Res. 34, 181–185.

Hartveit, E. (1999). Reciprocal synaptic interactions between rod bipolar cells and amacrine cells in the rat retina. J. Neurophysiol. 81, 2923–2936.

Hirai, H., Pang, Z., Bao, D., Miyazaki, T., Li, L., Miura, E., Parris, J., Rong, Y., Watanabe, M., Yuzaki, M., et al. (2005). Cbln1 is essential for synaptic integrity and plasticity in the cerebellum. Nat. Neurosci. 8, 1534–1541.

Ibata, K., Kono, M., Narumi, S., Motohashi, J., Kakegawa, W., Kohda, K., and Yuzaki, M. (2019). Activity-Dependent Secretion of Synaptic Organizer Cbln1 from Lysosomes in Granule Cell Axons. Neuron 102, 1184–1198.

Ichtchenko, K., Hata, Y., Nguyen, T., Ullrich, B., Missler, M., Moomaw, C., and Südhof, T.C. (1995). Neuroligin 1: A splice site-specific ligand for β-neurexins. Cell 81, 435–443.

Ichtchenko, K., Nguyen, T., and Südhof, T.C. (1996). Structures, alternative splicing, and neurexin binding of multiple neuroligins. J. Biol. Chem. 271, 2676–2682.

Isaacson, J.S. (2001). Mechanisms governing dendritic gamma-aminobutyric acid (GABA) release in the rat olfactory bulb. Proc Natl Acad Sci U S A 98, 337–342.

Isaacson, J.S., and Strowbridge, B.W. (1998). Olfactory reciprocal synapse: dendritic signalling in the CNS. Neuron 20, 749–761.

Ito-Ishida, A., Kakegawa, W., Kohda, K., Miura, E., Okabe, S., and Yuzaki, M. (2014). Cbln1 downregulates the formation and function of inhibitory synapses in mouse cerebellar Purkinje cells. Eur. J. Neurosci. 39, 1268–1280.

Ito-Ishida, A., Miyazaki, T., Miura, E., Matsuda, K., Watanabe, M., Yuzaki, M., and Okabe, S. (2012). Presynaptically Released Cbln1 Induces Dynamic Axonal Structural Changes by Interacting with GluD2 during Cerebellar Synapse Formation. Neuron 76, 549–564.

Jahr, C.E., and Nicoll, R.A. (1980). Dendrodendritic inhibition: Demonstration with intracellular recording. Science 207, 1473–1475.

Joo, J.Y., Lee, S.J., Uemura, T., Yoshida, T., Yasumura, M., Watanabe, M., and Mishina, M. (2011). Differential interactions of cerebellin precursor protein (Cbln) subtypes and neurexin variants for synapse formation of cortical neurons. Biochem. Biophys. Res. Commun. 406, 627–632.

Kasem, E., Kurihara, T., and Tabuchi, K. (2018). Neurexins and neuropsychiatric disorders. Neurosci. Res. 127, 53–60.

Knott, G.W., Quairiaux, C., Genoud, C., and Welker, E. (2002). Formation of dendritic spines with GABAergic synapses induced by whisker stimulation in adult mice. Neuron 34, 265–273.

Liu, Z., Chen, Z., Shang, C., Yan, F., Shi, Y., Zhang, J., Qu, B., Han, H., Wang, Y., Li, D., et al. (2017). IGF1-Dependent Synaptic Plasticity of Mitral Cells in Olfactory Memory during Social Learning. Neuron 95, 106–122.

Madisen, L., Zwingman, T.A., Sunkin, S.M., Oh, S.W., Zariwala, H.A., Gu, H., Ng, L.L., Palmiter, R.D., Hawrylycz, M.J., Jones, A.R., et al. (2010). A robust and high-throughput Cre reporting and characterization system for the whole mouse brain. Nat. Neurosci. 13, 133–140.

Matsuda, K., and Yuzaki, M. (2011). Cbln family proteins promote synapse formation by regulating distinct neurexin signaling pathways in various brain regions. Eur. J. Neurosci. 33, 1447–1461.

Matsuda, K., Miura, E., Miyazaki, T., Kakegawa, W., Emi, K., Narumi, S., Fukazawa, Y., Ito-Ishida, A., Kondo, T., Shigemoto, R., et al. (2010). Cbln1 Is a Ligand for an Orphan Glutamate Receptor d2, a Bidirectional Synapse Organizer. Science 338, 1541–1545.

Naritsuka, H., Sakai, K., Hashikawa, T., and Mori, K. (2009). Perisomatic-Targeting Granule Cells in the Mouse. J. Comp. Neurol. 426, 409–426.

Nguyen, T., and Südhof, T.C. (1997). Binding properties of neuroligin 1 and neurexin 1β reveal function as heterophilic cell adhesion molecules. J. Biol. Chem. 272, 26032–26039.

Pang, Z., Zuo, J., and Morgan, J.I. (2000). Cbln3, a novel member of the precerebellin family that binds specifically to Cbln1. J. Neurosci. 20, 6333–6339.

Pressler, R.T., and Strowbridge, B.W. (2017). Direct Recording of Dendrodendritic Excitation in the Olfactory Bulb: Divergent Properties of Local and External Glutamatergic Inputs Govern Synaptic Integration in Granule Cells. J. Neurosci. 37, 11774–11788.

Rall, W., Shepherd, G.M., Reese, T.S., and Brightman, M.W. (1966). Dendrodendritic synaptic pathway for inhibition in the olfactory bulb. Exp. Neurol. 14, 44–56.

Reissner, C., Klose, M., Fairless, R., and Missler, M. (2008). Mutational analysis of the neurexin/neuroligin complex reveals essential and regulatory components. Proc. Natl. Acad. Sci. U. S. A. 105, 15124–15129.

Sanz, E., Yang, L., Su, T., Morris, D.R., McKnight, G.S., and Amieux, P.S. (2009). Cell-type-specific isolation of ribosome-associated mRNA from complex tissues. Proc. Natl. Acad. Sci. U. S. A. 106, 13939–13944.

Schneider, C.A., Rasband, W.S., and Eliceiri, K.W. (2012). NIH Image to ImageJ: 25 years of image analysis. Nat. Methods 9, 671–675.

Seigneur, E., and Südhof, T.C. (2018). Genetic ablation of all cerebellins reveals synapse organizer functions in multiple regions throughout the brain. J. Neurosci. 38, 4774–4790.

Shepherd, G.M., Hines, M.L., Migliore, M., Chen, W.R., and Greer, C.A. (2020). Predicting brain organization with a computational model: 50-year perspective on lateral inhibition and oscillatory gating by dendrodendritic synapses. J. Neurophysiol. 124, 375–387.

Sloper, J.J.; Powell, T.P.S. (1978). Dendro-Dendritic and Reciprocal Synapses in the Primate Motor Cortex. Proc. R. Soc. London. Ser. B. Biol. Sci. 203, 23–38.

Sterky, F.H., Trotter, J.H., Lee, S., Recktenwald, C. V, Du, X., Zhou, B., Zhou, P., Schwenk, J., Fakler, B., and Südhof, T.C. (2017). Carbonic anhydrase-related protein CA10 is an evolutionarily conserved pan-neurexin ligand. Proc. Natl. Acad. Sci. U. S. A. 114, E1253–1262.

Südhof, T.C. (2017). Synaptic Neurexin Complexes: A Molecular Code for the Logic of Neural Circuits. Cell 171, 745–769.

Südhof, T.C. (2021). The Cell Biology of Synapse formation. J. of Cell Biology, in press.

Tabuchi, K., and Südhof, T.C. (2002). Structure and evolution of neurexin genes: Insight into the mechanism of alternative splicing. Genomics 79, 849–859.

Tervo, D.G.R., Hwang, B.-Y., Viswanathan, S., Gaj, T., Lavzin, M., Ritola, K., Lindo, S., Michael, S., Kuleshova, E., Ojala, D., et al. (2016). A Designer AAV Variant Permits Efficient Retrograde Access to Projection Neurons. Neuron 92, 372–382.

Trotter, J.H., Dargaei, Z., Sclip, A., Essayan-Perez, S., Liakath-Ali, K., Raju, K., Nabet, A., Liu, X., and Südhof, T.C. (2020). Compartment-Specific Neurexin Nanodomains Orchestrate Tripartite Synapse Assembly. BioRxiv 2020.08.21.262097.

Uchigashima, M., Cheung, A., Suh, J., Watanabe, M., and Futai, K. (2019). Differential expression of neurexin genes in the mouse brain. J. Comp. Neurol. 527, 1940–1965.

Uemura, T., Lee, S.J., Yasumura, M., Takeuchi, T., Yoshida, T., Ra, M., Taguchi, R., Sakimura, K., and Mishina, M. (2010). Trans-synaptic interaction of GluRδ2 and Neurexin through Cbln1 mediates synapse formation in the cerebellum. Cell 141, 1068–1079.

Ullrich, B., Ushkaryov, Y.A., and Südhof, T.C. (1995). Cartography of neurexins: More than 1000 isoforms generated by alternative splicing and expressed in distinct subsets of neurons. Neuron 14, 497–507.

Urban, N.N., and Arevian, A.C. (2009). Computing with dendrodendritic synapses in the olfactory bulb. Ann. N. Y. Acad. Sci. 1170, 264–269.

Ushkaryov, Y.A., and Südhof, T.C. (1993). Neurexin IIIα: Extensive alternative splicing generates membrane-bound and soluble forms. Proc. Natl. Acad. Sci. U. S. A. 90, 6410–6414.

Ushkaryov, Y.A., Petrenko, A.G., Geppert, M., and Südhof, T.C. (1992). Neurexins: Synaptic cell surface proteins related to the α-latrotoxin receptor and laminin. Science 257, 50–56.

Varoqueaux, F., Jamain, S., and Brose, N. (2004). Neuroligin 2 is exclusively localized to inhibitory synapses. Eur. J. Cell Biol. 83, 449–456.

Wang, C.Y., Liu, Z., Ng, Y.H., and Südhof, T.C. (2020). A Synaptic Circuit Required for Acquisition but Not Recall of Social Transmission of Food Preference. Neuron 107, 144–157.

Yasumura, M., Yoshida, T., Lee, S.J., Uemura, T., Joo, J.Y., and Mishina, M. (2012). Glutamate receptor δ1 induces preferentially inhibitory presynaptic differentiation of cortical neurons by interacting with neurexins through cerebellin precursor protein subtypes. J. Neurochem. 121, 705–716.

Yuzaki, M. (2018). Two Classes of Secreted Synaptic Organizers in the Central Nervous System. Annu. Rev. Physiol. 80, 243–262.

Zhang, B., and Südhof, T.C. (2016). Neuroligins are selectively essential for NMDAR signaling in cerebellar stellate interneurons. J. Neurosci. 36, 9070–9083.

Zhang, B., Chen, L.Y., Liu, X., Maxeiner, S., Lee, S.J., Gokce, O., and Südhof, T.C. (2015). Neuroligins Sculpt Cerebellar Purkinje-Cell Circuits by Differential Control of Distinct Classes of Synapses. Neuron 87, 781–796.

Zhang, B., Gokce, O., Hale, W.D., Brose, N., and Südhof, T.C. (2018). Autism-associated neuroligin-4 mutation selectively impairs glycinergic synaptic transmission in mouse brainstem synapses. J. Exp. Med. 215, 1543–1553.

Zhou, F.W., Shao, Z.Y., Shipley, M.T., and Puche, A.C. (2020). Short-term plasticity in glomerular inhibitory circuits shapes olfactory bulb output. J. Neurophysiol. 123, 1120–1132.

