## Supplementary figures for "The Molecular Logic Organizing the Functional Compartmentalization of Reciprocal Synapses"

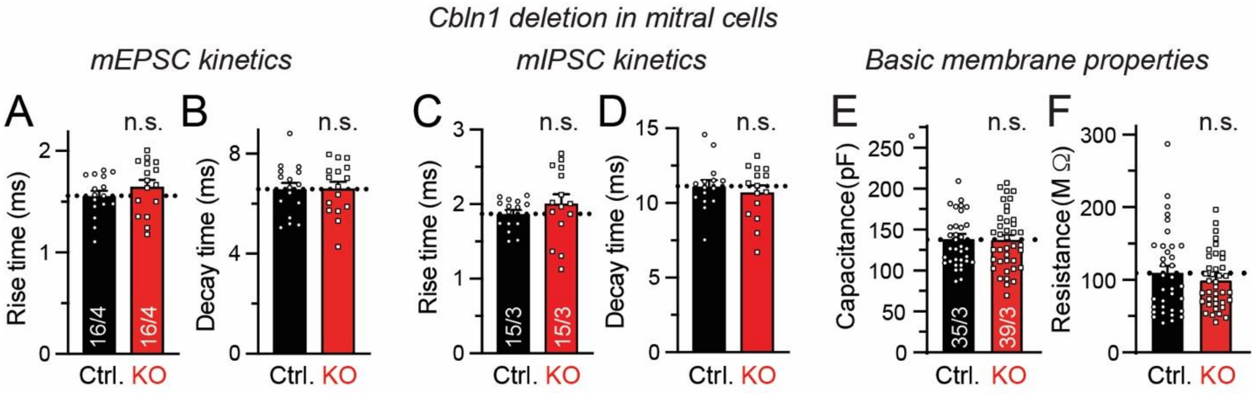


**Supplementary Figure 1. Analyses of the kinetics of mEPSCs and mIPSCs and of basic membrane properties of mitral cells do not reveal any changes induced by the postsynaptic *Cbln1* deletion (related to Fig. 2).**

***A***. mEPSC rise time (10% to 90%).

***B***. mEPSC decay time (90% to 10%).

***C***. mIPSC rise time (10% to 90%).

***D***. mIPSC decay time (90% to 10%).

***E***. Membrane capacitance.

***F***. Membrane resistance.

All numerical data are means ± SEM (number of cells/experiments examined is listed in the bars of A, C and E). Statistical analyses performed by *Student's* t-test uncovered no statistically significant differences.

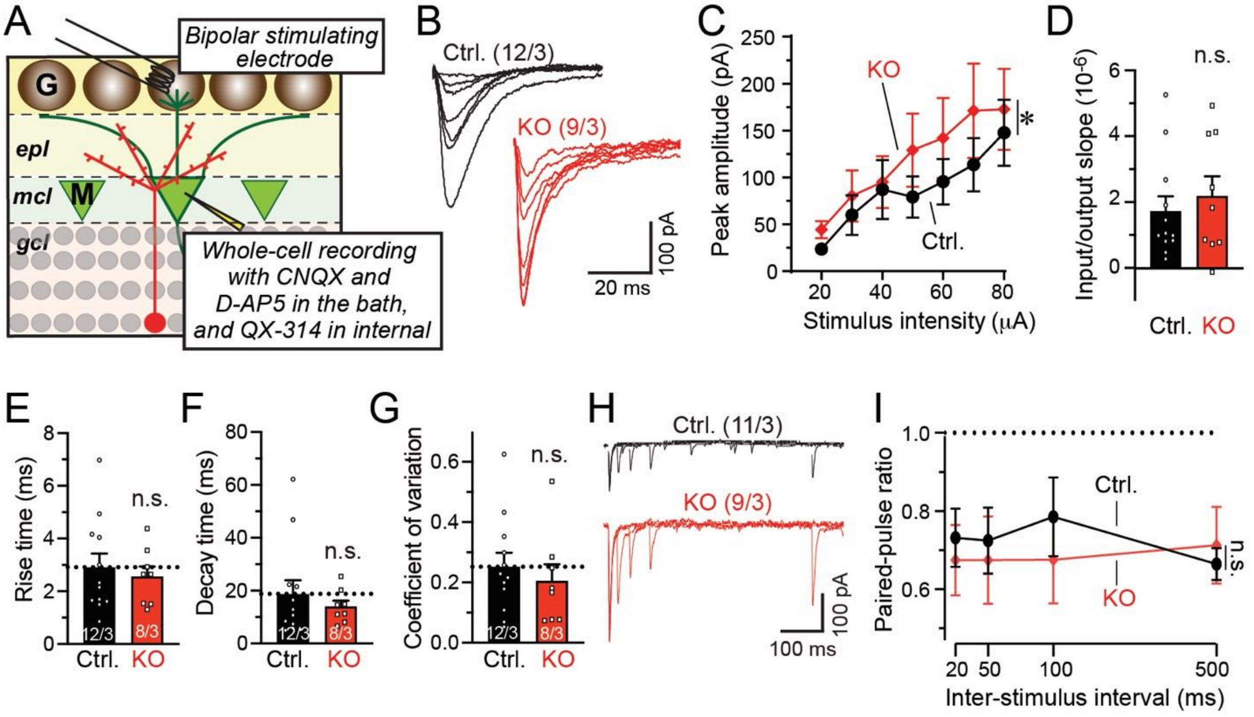


**Supplementary Figure 2. Periglomerular cell (PGC)-to-mitral cell (MC) evoked IPSCs are not altered by the *Cbln1* deletion from mitral cells (related to Fig. 3).**

***A***. Recording paradigm.

***B***. Representative traces of evoked PGC→MC IPSCs.

***C & D***. Input/output curve of the PGC→MC IPSC peak amplitude (C) and summary of its slope (D).

***E & F***. Kinetics of PGC→MC IPSCs (E, rise time; F, decay time; both measured at a 70 µA stimulus intensity).

***G***. Coefficient of variations of evoked IPSC at a 70 µA stimulus intensity.

***H & I***. Representative traces (H) and summary plot (I) of the paired-pulse ratios of evoked IPSCs measured at 70 µA stimulus intensity with different interstimulus intervals.

All numerical data are means ± SEM (numbers of cells/mice analyzed are indicated above the sample traces and bars in E-G). Statistical analyses were performed by by *Student's* t-test in D-G, and by two-way ANOVA with Bonferroni's multiple comparison test in C and I (*, $p<0.05$). One cell in KO groups was excluded from kinetics and CV analysis because its peak amplitude was close to zero even at maximal stimulus, which would skew kinetics and CV analysis.

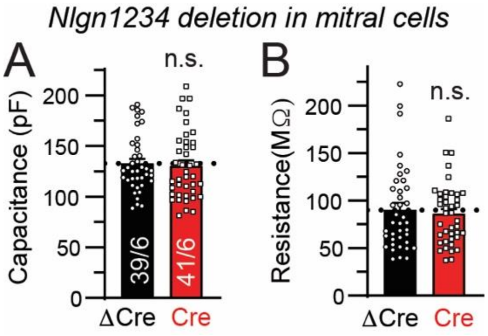


**Supplementary Figure 3. Basic membrane properties as a function of *Nlgn1234* deletion in mitral cells (related to Fig. 5)**

***A***. Membrane capacitance;

***B***. Membrane resistance.

All numerical data are means ± SEM (number of cells/experiments examined is shown in the bars of A). Statistical analyses performed by *Student*'s t-test uncovered no statistically significant differences.

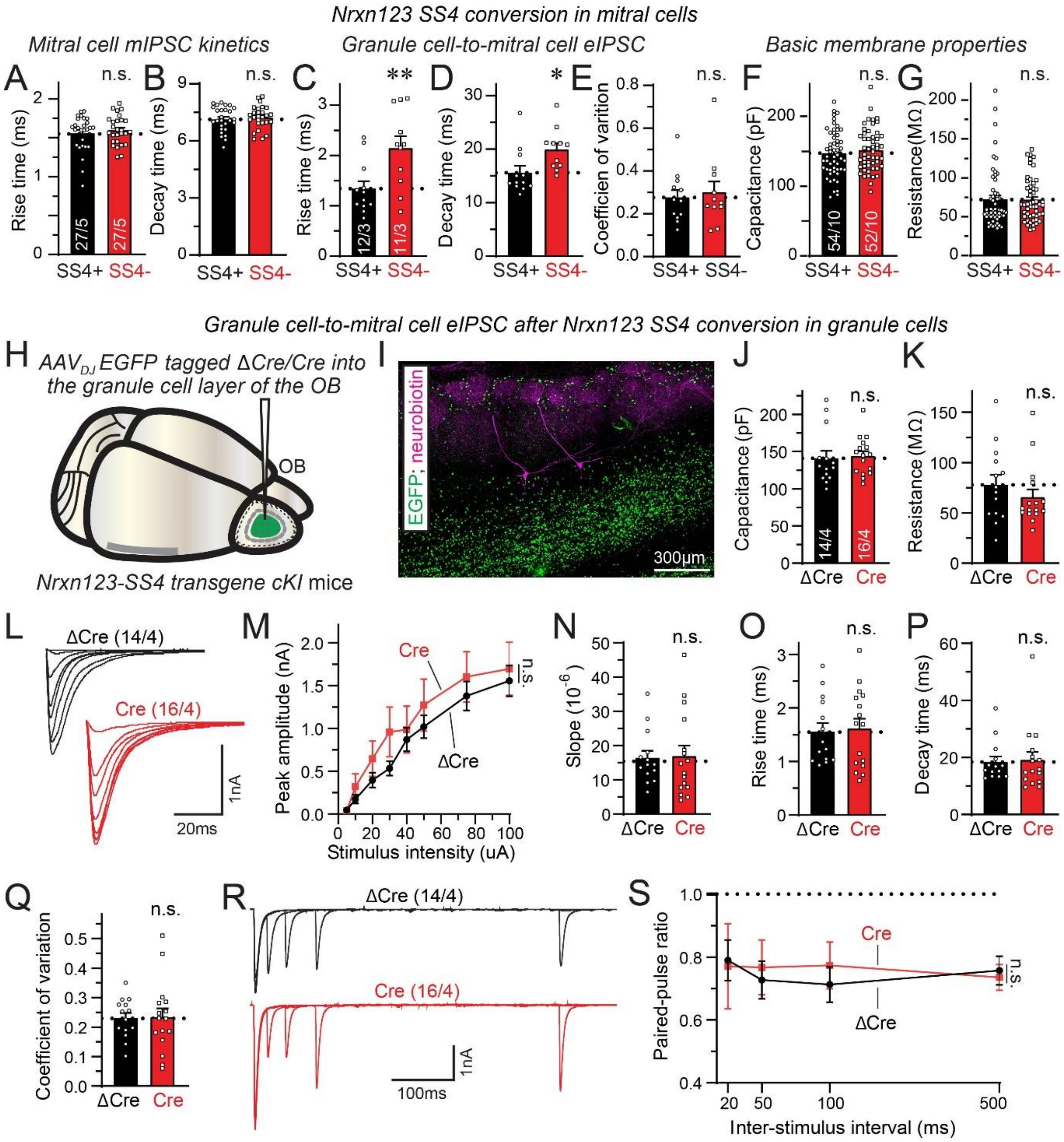


**Supplementary Figure 4. Additional electrophysiological measurements that analyze the effect of SS4+ vs. SS4- variants of all neurexins using *Nrxn123*-SS4+ conditional knockin mice (related to Fig. 6)**. All experiments as well as those shown in Figure 6 were performed with conditional knockin mice in which all neurexin genes were mutated to constitutively express all neurexins as SS4+ variants, but in which these SS4+ neurexins can be converted into SS4- neurexins by Cre recombination of the mutant genes (Dai et al., 2019).

***A-G***. Additional electrophysiological measurements of mitral cells as a function of the SS4+ vs. SS4- variant of all neurexins. Note that there is no difference in mitral mIPSC kinetics (A, mIPSC rise times (10% to 90%); B, mIPSC decay times (90% to 10%)), but the the rise and decay times of evoked IPSCs are elevated (C, GC→MC IPSC rise times; D, GC→MC IPSC decay times), while the coefficient of variation of evoked GC→MC IPSCs is unchanged (E), and the intrinsic membrane properties are also not altered (F, membrane capacitance; G, membrane resistance).

***H***. Experimental strategy to convert SS4+ into SS4- variants of all neurexins in granule cells of the OB in *Nrxn123*-SS4+ conditional KI mice. When AAVs encoding Cre or ΔCre (as a control) are stereotactically injected directly into the OB, the AAVs almost exclusively infect granule cells and spare mitral cells.

***I***. Representative image of an acute slice from an OB that was infected in vivo with AAVs (green fluorescence is due to EGFP-tagged Cre or ΔCre). Two mitral cells in the slice were patched and filled neurobiotin (magenta).

***J & K***. Passive membrane properties of mitral cells as a function of granule-cell *Nrxn123* SS4 conversion (J, membrane capacitance; K, membrane resistance).

***L-S***. Conversion of SS4+ neurexins into SS4- neurexins in granule cells has no effect on the properties and strength of GC→MC synaptic transmission (L, representative traces of GC→MC IPSCs; M & N, input/output curve of the peak amplitude of evoked GC→MC IPSCs (M) and summary of the input/output curve slope (N); O & P: kinetics of GC→MC IPSCs (O, rise times; P, decay times; both measured at a 75 µA stimulus intensity); Q, coefficient of variation of evoked IPSCs at a 75 µA stimulus intensity; R & S, representative traces (R) and summary plot (S) of the paired-pulse ratio of evoked IPSCs measured at a 75 µA stimulus intensity).

All numerical data are means ± SEM (number of cells/experiments examined is shown in the bars of A, D, F, and J, or above the sample traces in L and R). Statistical analyses performed by two-way ANOVA (M and S) or *Student's* t-test (all other data) uncovered no statistically significant differences except in C and D (*: $p<0.05$; **: $p<0.01$).

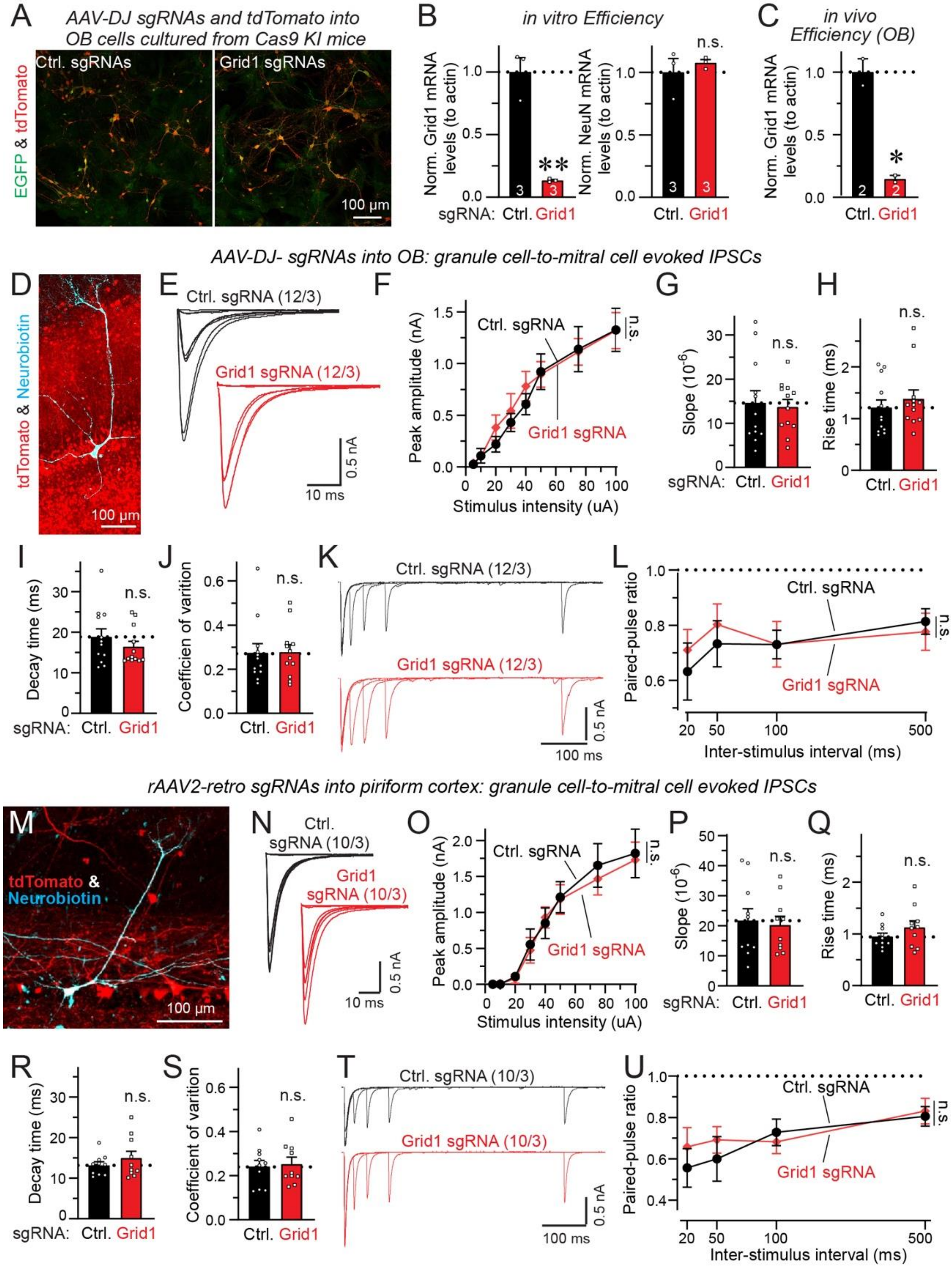
A
AAV-DJ sgRNAs and tdTomato into OB cells cultured from Cas9 KI mice
Ctrl. sgRNAs
Grid1 sgRNAs
EGFP & tdTomato
100 µm
B
in vitro Efficiency
Norm. Grid1 mRNA levels (to actin)
Norm. NeuN mRNA levels (to actin)
n.s.
sgRNA: Ctrl. Grid1
C
in vivo Efficiency (OB)
Norm. Grid1 mRNA levels (to actin)
Ctrl. Grid1
AAV-DJ- sgRNAs into OB: granule cell-to-mitral cell evoked IPSCs
D
tdTomato & Neurobiotin
100 µm
E
Ctrl. sgRNA (12/3)
Grid1 sgRNA (12/3)
0.5 nA
10 ms
F
Peak amplitude (nA)
Ctrl. sgRNA
Grid1 sgRNA
Stimulus intensity (uA)
n.s.
G
Slope (10^-6)
n.s.
sgRNA: Ctrl. Grid1
H
Rise time (ms)
n.s.
Ctrl. Grid1
I
Decay time (ms)
n.s.
sgRNA: Ctrl. Grid1
J
Coefficien of varition
n.s.
Ctrl. Grid1
K
Ctrl. sgRNA (12/3)
Grid1 sgRNA (12/3)
0.5 nA
100 ms
L
Paired-pulse ratio
Ctrl. sgRNA
Grid1 sgRNA
n.s.
Inter-stimulus interval (ms)
rAAV2-retro sgRNAs into piriform cortex: granule cell-to-mitral cell evoked IPSCs
M
tdTomato & Neurobiotin
100 µm
N
Ctrl. sgRNA (10/3)
Grid1 sgRNA (10/3)
0.5 nA
10 ms
O
Peak amplitude (nA)
Ctrl. sgRNA
Grid1 sgRNA
n.s.
Stimulus intensity (uA)
P
Slope (10^-6)
n.s.
sgRNA: Ctrl. Grid1
Q
Rise time (ms)
n.s.
Ctrl. Grid1
R
Decay time (ms)
n.s.
sgRNA: Ctrl. Grid1
S
Coefficient of varition
n.s.
Ctrl. Grid1
T
Ctrl. sgRNA (10/3)
Grid1 sgRNA (10/3)
0.5 nA
100 ms
U
Paired-pulse ratio
Ctrl. sgRNA
Grid1 sgRNA
n.s.
Inter-stimulus interval (ms)

**Supplementary Figure 5. The effect of *Cbln1* deletion in mitral/tufted cells on GC→MC synaptic transmission does not depend on mitral cell GluD1 signaling (related to Fig. 6)**

***A***. Representative images of transfected cells cultured from the OB of newborn wildtype Cas9 KI mice. Cells were infected with $AAV_{DJ}$ co-expressing control or Grid1-targeting sgRNAs and tdTomato (Dai et al., 2021).

***B***. *in vitro* efficiency of CRISPR/Cas9-mediated deletion of GluD1 measured by quantitative RT-PCR (left, GluD1 mRNA levels; right, NeuN mRNA levels).

***C***. *in vivo* efficiency of CRISPR/Cas9-mediated deletion of GluD1 measured by RT-qPCR. mRNA was extracted from olfactory bulbs injected with $AAV_{DJ}$ co-expressing control or Grid1 sgRNAs and tdTomato.

***D-L***. GC→MC evoked IPSCs as a function of the global *Grid1* deletion in the OB (D, representative images of injected olfactory bulb and recorded mitral cell; E: representative traces of GC→MC evoked IPSCs; F & G, input/output curve of GC→MC evoked IPSCs peak amplitude (F) and summary of its slope (G); H & I, kinetics of GC→MC evoked IPSCs (H: rise time; I: decay time; both measured at 75µA stimulus intensity); J, coefficient of variations of evoked IPSCs at a 75 µA stimulus intensity; K & L, representative traces (K) and summary plot (L) of paired-pulse ratio of evoked IPSCs measured at a 75 µA stimulus intensity.

***M-U***. GC→MC evoked IPSCs as a function of mitral-cell specific *Grid1* deletion through rAAV2-retro injection into piriform cortex, with a panel arrangement similar to that of D-L.

All numerical data are means ± SEM. Number of culture wells are indicated in bars in B, number of mice are indicated in bars in C, and number of cells/mice analyzed are indicated above the sample traces in D-L. Statistical analyses were performed by *Student's* t-test in B, C, G-J and P-S, and by two-way ANOVA with Bonferroni's multiple comparison test in F, L, O and U (*, $p<0.05$; **, $p<0.01$).

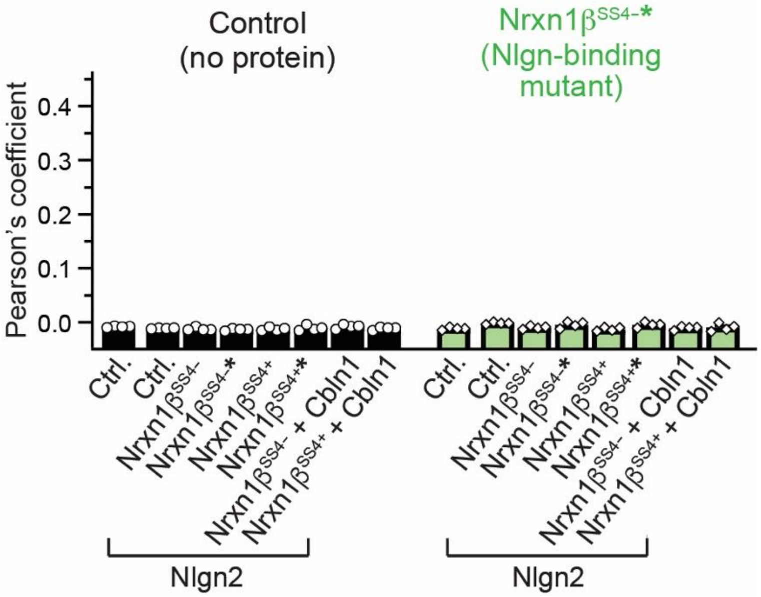


**Supplementary Figure 6. Summary graph of cell aggregation mediated by control groups in the *cis*- vs. *trans*-interaction assay of Figure 7**

Summary of Pearson's coefficient related to control (empty vectors) and Nrxn1β$^{SS4-}$* (Nlgn-binding mutant) groups. Data are means ± SEM (n=4 batches of culture).

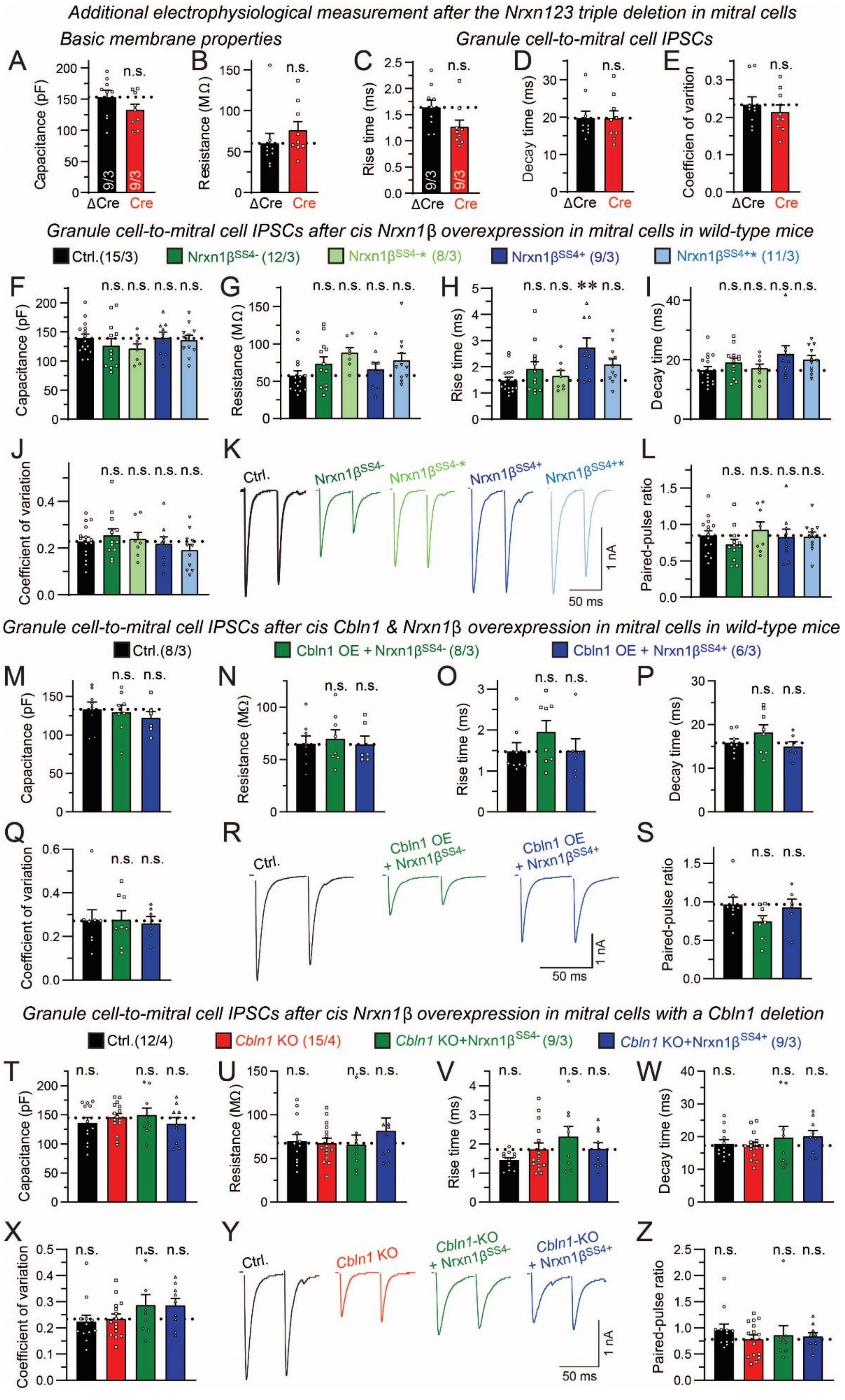
Additional electrophysiological measurement after the Nrxn123 triple deletion in mitral cells
Basic membrane properties
Granule cell-to-mitral cell IPSCs
Granule cell-to-mitral cell IPSCs after cis Nrxn1β overexpression in mitral cells in wild-type mice
Granule cell-to-mitral cell IPSCs after cis Cbln1 & Nrxn1β overexpression in mitral cells in wild-type mice
Granule cell-to-mitral cell IPSCs after cis Nrxn1β overexpression in mitral cells with a Cbln1 deletion

**Supplementary Figure 7. Additional electrophysiological data obtain in mitral cells after deletion of all neurexins in *Nrxn123* conditional triple KO mice (A-E) or after overexpression of various Nrxn1β (or together with Cbln1) proteins in wild-type or *Cbln1* KO cells (F-Z) (related to Fig. 8).**

***A*-*E***. Additional electrophysiological measurements in mitral cells as a function of the mitral cell-specific *Nrxn123* deletion (A, membrane capacitance; B, membrane resistance; C & D, rise and decay times, respectively, of evoked GC→MC IPSCs (measured at 75 µA stimulus intensity); E, coefficient of variation of GC→MC IPSCs at a 75 µA stimulus intensity).

***F*-*L***. Additional electrophysiological measurements in mitral cells as a function of the mitral cell-specific overexpression of various Nrxn1β proteins in wild-type mice (F, membrane capacitance; G, membrane resistance. H & I, rise and decay times, respectively, of evoked GC→MC IPSCs (measured at 75 µA stimulus intensity); J, coefficient of variation of IPSCs at a 75 µA stimulus intensity; K & L, representative traces (K) and summary plot (L) of paired-pulse ratio measurements of evoked IPSCs monitored at a 75 µA stimulus intensity.

***M*-*S***. Additional electrophysiological measurements in mitral cells as a function of the mitral cell-specific co-overexpression of Cbln1 and Nrxn1β$^{SS4+}$ (or Nrxn1β$^{SS4-}$) in wild-type mice. Cbln1 overexpression (OE) was achieved by rAAV2-retro expressing Cbln1-P2A-EGFP, and EGFP/tdTomato co-expression was used in the control group. Arrangements of panels is the same as in F-L.

***T*-*Z***. Additional electrophysiological measurements in mitral cells as a function of the mitral cell-specific overexpression of Nrxn1β$^{SS4+}$ and Nrxn1β$^{SS4-}$ in *Cbln1* KO mice. *Cbln1 cKO* mice without (Ctrl.) or with a *tBet-Cre* allele (*Cbln1* KO) were also infected retrogradely in mitral cells with *Nrxn1β*-overexpressing rAAV2-retros. Arrangements of panels is the same as in F-L.

All numerical data are means ± SEM (number of cells/experiments examined is shown in the bars of A and C, or in the assignment bars of colors to experimental conditions). Statistical analyses performed by Student's t-test (A-E) or one-way ANOVA with Bonferroni's multiple comparison test (with comparison to the Ctrl. in F-L and M-S and to the *Cbln1* KO group in T-Z) uncovered no statistically significant differences except in H (**: $p<0.01$).
